# Drivers of population dynamics of at-risk populations change with pathogen arrival

**DOI:** 10.1101/2023.11.27.568870

**Authors:** Alexander T. Grimaudo, Joseph R. Hoyt, R. Andrew King, Rickard S. Toomey, Chris Simpson, Cory Holliday, Alexander Silvis, Rick T. Doyle, Joseph A. Kath, Mike P. Armstrong, Virgil Brack, Richard J. Reynolds, Ryan H. Williamson, Gregory G. Turner, Vona Kuczynska, Jordan J. Meyer, Kyle Jansky, Carl J. Herzog, Skylar R. Hopkins, Kate E. Langwig

## Abstract

Successful wildlife conservation in an era of rapid global change requires understanding determinants of species population abundance and growth. However, when populations are faced with novel stressors, factors associated with healthy and growing populations can change, necessitating a shift in conservation strategies. For example, emerging infectious diseases can cause conditions previously beneficial or neutral to host populations to increase disease impacts. Here, we paired a population dataset of 265 colonies of the federally endangered Indiana bat (*Myotis sodalis*) with 50.7 logger-years of environmental data to explore factors that affected colony response to white-nose syndrome (WNS), an emerging fungal disease. We found wide variation in colony responses to WNS, ranging from extirpation to stabilization and persistence. Simulating future population dynamics suggests that most extirpations have already occurred, as the pathogen has been present for several years in most colonies, and that small colonies were more susceptible to extirpation than large ones. Further, while temperature and humidity conditions of hibernacula appeared unassociated with Indiana bat colony growth prior to WNS, extirpation risk following pathogen arrival was elevated in colonies that used colder and wetter hibernacula. Additionally, rates of decline were greater in colder hibernacula, opposite the association for a sympatric bat species. Overall, this study illustrates that emerging infectious diseases can change the factors associated with host population abundance and optimal growth, including through novel environmental associations, which can vary across host species. Consideration of these shifting associations and intrinsic differences between impacted host species will be essential to successful species conservation.

## Introduction

Conserving global biodiversity in the Anthropocene requires a deep understanding of the factors that contribute to the resilience and growth of wildlife populations. The suite of biotic and abiotic conditions that can contribute to population growth, from ambient conditions like temperature (Woodworth et al., 2017), humidity (Lu and Wu, 2011), or precipitation (Masó et al., 2020), to community-level factors like predator density (Salo et al., 2010), competitor density (Gamelon et al., 2019), or pathogen prevalence (Hoyt et al., 2020), can further serve as targets for conservation and management. However, novel stressors such as climate change (Acevedo et al., 2020; Şekercioĝlu et al., 2012; Van Dyck et al., 2015), invasive species introduction (Hagman et al., 2009), and habitat fragmentation and loss (Haddad et al., 2015; Mantyka-Pringle et al., 2015) can alter associations between biotic and abiotic conditions and population growth or stability. The form and magnitude of these altered associations is often ambiguous or difficult to characterize and can create the potential for a mismatch between conservation interventions and intended outcomes, possibly with negative effects on the targeted population (Sutherland et al., 2004). Therefore, understanding how novel stressors change the way wildlife populations respond to biotic and abiotic conditions will be essential to their successful conservation in a rapidly changing world.

Emerging infectious diseases are one of many stressors that threaten wildlife populations and can result in host species declines or population extirpations (Altizer et al., 2013; Anderson et al., 2004; Daszak et al., 2001, 2000; Fisher et al., 2012; Rogalski et al., 2017), as evidenced by West Nile Virus and avian malaria in birds (Kilpatrick, 2011; Van Riper III et al., 1986), chytridiomycosis in amphibians (Skerratt et al., 2007), snake fungal disease (Lorch et al., 2016), sarcoptic mange in wombats (Martin et al., 2018), facial tumor disease in Tasmanian devils (Mccallum, 2008), and white-nose syndrome in bats (Hoyt et al., 2021). Early conservation measures to bolster the demographic resilience and adaptability of host populations might dampen the impact of disease following its arrival (Grant et al., 2017; Hodgson et al., 2015). However, it is often unclear whether those same conservation measures will promote the persistence of the host population during pathogen invasion when pathogen transmission dynamics (Huang et al., 2021; Leach et al., 2016) or environmental dependence of disease (Langwig et al., 2012; McNew et al., 2019; Samuel et al., 2015) can significantly shift the fitness landscape of the host. This information is particularly important for rare and endangered host species with limited habitat availability or population sizes before pathogen invasion, as their sensitivity to novel stressors and management interventions necessitates unequivocal conservation best practices.

The size of a host population can contribute to its response to pathogen invasion and ultimately its risk of extirpation. Greater standing genetic diversity in large host populations may promote evolutionary adaptation to an invading pathogen before total population collapse occurs (Barrett and Schluter, 2008; Bell, 2013; Carlson et al., 2014; Gomulkiewicz and Holt, 1995; Maslo and Fefferman, 2015). Large populations also have less risk of stochastic fadeout, which can be exacerbated in populations declining from disease (de Castro and Bolker, 2005; Frick et al., 2015; Langwig et al., 2012). However, if large populations also have high host densities, pathogen transmission may be elevated, leading to more rapid declines and greater extinction risk (Brunner et al., 2017; Cross et al., 2010; Cunningham et al., 2021; Rachowicz and Briggs, 2007; Smith et al., 2009; Storm et al., 2013). While there are clear theoretical mechanisms of how host population size can contribute to its response to an emerging infectious disease, this association is often left uncharacterized due to the difficulty of obtaining sufficient data, despite being essential to prioritizing populations for conservation action.

Environmental dependence of pathogen transmission or disease severity can create both hotspots (Lambin et al., 2010; Ostfeld et al., 2005; Paull et al., 2012) or refugia from (Heard et al., 2015; Mosher et al., 2018; Puschendorf et al., 2011; Scheele et al., 2017; Spitzen-Van Der Sluijs et al., 2017) infection and disease. Temperature, for example, can have complex effects on disease by influencing pathogen growth rates (Hopkins et al., 2021), pathogen transmission (Mordecai et al., 2019), host susceptibility to infection (Cohen et al., 2017), or disease severity following infection (Debes et al., 2017), processes that scale up to variation in population-level impacts on the host. These environmental effects can further exacerbate impacts to host populations if high-quality habitat prior to pathogen invasion becomes population sinks or ecological traps of high pathogen transmission and disease severity following disease emergence (Hopkins et al., 2021; Leach et al., 2016). Understanding environmental contributions to host population response offers a point of leverage for conservation efforts, as protection and enhancement of critical environmental conditions is fundamental to the conservation of threatened host species. Importantly, environmental associations need not take the same form both pre- and post-pathogen arrival, and effective approaches to host species conservation must incorporate these dynamic processes.

White-nose syndrome (WNS) is an emerging infectious disease of hibernating bats caused by the fungal pathogen *Pseudogymnoascus destructans* (Lorch et al., 2011; Minnis and Lindner, 2013; Warnecke et al., 2012). The pathogen was introduced to North America in the early 2000s from its origin in Eurasia and has since spread widely through naïve hibernating bat populations (Blehert et al., 2009; Drees et al., 2017; Leopardi et al., 2015), causing severe population declines (Frick et al., 2015; Hoyt et al., 2021; Langwig et al., 2016, 2012). *Pseudogymnoascus destructans* establishes an environmental pathogen reservoir within winter hibernacula (Hoyt et al., 2020, 2015; Laggan et al., 2023), the caves and mines where bats hibernate, and can persist for years in the absence of its host (Hoyt et al., 2015). This results in seasonal infection dynamics in which WNS prevalence increases as bats arrive in hibernacula in late fall but declines during spring and summer months when bats do not primarily use hibernacula for roosting and bats are often euthermic (Langwig et al., 2015). Infection severity increases over the winter hibernation period when body temperatures drop to approximately ambient conditions (Langwig et al., 2017, 2015), allowing the cold-adapted fungus to invade the epidermal tissue (Lorch et al., 2011; Warnecke et al., 2013). Infection with *P. destructans* is accompanied by a suite of physiological changes that lead to dehydration (Cryan et al., 2013; Mcguire et al., 2017; Verant et al., 2014), frequent arousals from torpor (Lilley et al., 2016; Reeder et al., 2012; Warnecke et al., 2012), emaciation, and ultimately mortality (Cryan et al., 2010; Warnecke et al., 2013).

The Indiana bat (*Myotis sodalis*) is a North American species that has experienced severe but variable population declines due to WNS (Langwig et al., 2012; Thogmartin et al., 2012). Despite its status as endangered under the U.S. Endangered Species Act, little is known about drivers of variation in Indiana bat declines and persistence with disease, information important for its conservation. Environmental correlates of disease severity and population response have been previously reported for other species, particularly the little brown bat (*Myotis lucifugus*), where warmer hibernacula are associated with severe disease and declines (Grieneisen et al., 2015; Grimaudo et al., 2022; Hopkins et al., 2021; Langwig et al., 2016, 2012; Lilley et al., 2018), likely due to enhanced pathogen growth under those conditions (Marroquin et al., 2017; Verant et al., 2012). However, previous work has not found a clear association between temperature conditions of hibernacula and the response of Indiana bat colonies (Langwig et al., 2012). Evidence additionally suggests that for many species impacted by WNS, colony size may be related to the initial population response to the disease (Frick et al., 2015; Langwig et al., 2012). However, the particularly gregarious nature and large cluster sizes of Indiana bats (Clawson et al., 1980; Hardin and Hassell, 1970; Thomson, 1982) may result in constant density across colony sizes and time, keeping pathogen transmission and declines high regardless of colony size.

In this study, we pair an extensive bat population dataset curated by the U.S. Fish and Wildlife Service with a hibernacula environmental dataset to identify potential contributions of host population size and environmental conditions to Indiana bat population trends prior, during, and after pathogen arrival. Our data provide information on how population-level processes and variation in environmental conditions can generate heterogeneous host population responses to novel infectious diseases, lending insights into how to successfully manage their impacts.

## Material and Methods

### Data collection

#### Population data

We analyzed data on population sizes of 265 colonies of hibernating Indiana bats from 15 U.S. states collected between 2003—2022 (Supplemental Table 1). Population data were collected by various federal and state agencies and were compiled by the U.S. Fish and Wildlife Service. For each colony, the year of WNS detection within their hibernaculum was recorded and used to assign ‘epidemic year,’ which is the year since WNS detection (e.g., epidemic year 0 corresponds to the year of WNS detection). We classified census data by epidemic phase, with the ‘Pre-Invasion’ phase being epidemic years less than or equal to 0, ‘Epidemic’ being epidemic years 1 through 5, and ‘Established’ being epidemic years 6 or greater. This classification reflects observed patterns in colony responses to WNS, in which major declines begin in the year following WNS detection (epidemic year 1) and continue for several years until population stability is achieved by epidemic year 6, on average (Hoyt et al., 2021). Overall, census data was available for 214 hibernacula during the ‘Pre-invasion’ phase, 251 hibernacula during the ‘Epidemic’ phase, and 106 hibernacula during the ‘Established’ phase.

We estimated the annual population growth rates, λ_i_, for each Indiana bat colony for each winter with an available colony census. λ_i_ was derived by dividing the census value taken in a winter to the next most recent winter census value, separated by T years:

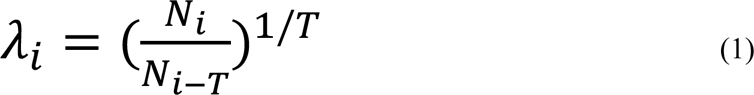

Mean population growth rates, λ_m_, for each colony in each epidemic phase were calculated by averaging all available annual lambda values for a colony within each epidemic phase. We omitted from the dataset any λ_i_ values that were calculated with N_i-T_ values less than 10, as calculations of λ_i_ are disproportionately sensitive at small colony sizes, producing misleadingly small or large λ_i_ values.

We assigned a classification to each colony based on their overall response to WNS, referred to as ‘colony status.’ ‘Extirpated’ colonies were those with a most recent post-WNS arrival (epidemic year greater than or equal to 0) census value of 0, indicating total collapse of the colony. ‘Declining’ colonies were those that never exhibited a year of stable or positive annual colony growth (λ_i_ greater than or equal to 0) after epidemic year 1. Finally, ‘persisting’ colonies were those that had at least one year of stable or positive annual colony growth after epidemic year 1. These classifications were possible for 231 of the 265 colonies in the population dataset. The remaining 34 colonies did not have sufficient post-WNS detection census data to accurately assign a classification, as they did not have census data available following epidemic year 0.

#### Environmental data

Data on the temperature and humidity conditions of Indiana bat roosting locations within hibernacula were collected overwinter in a total of 48 hibernacula between 1994 and 2022, 53.9% of which was collected since 2015 and 10.8% of which was collected as a part of this study. In total, the environmental dataset we used in this study constitutes 50.7 logger-years of data on the environmental conditions within hibernacula (summarized in Supplemental Table 1). Environmental data were recorded by a variety of data loggers, including Onset HOBO H8 Pro, Onset HOBO Pro V2, Onset HOBO Pendant, Onset HOBO XT, Madgetech TransiTempII-RH, Maxim Integrade iButton, and modified onset HOBO MX2303 data loggers (Supplemental Table 2). HOBO MX2303 units were modified into wet bulb psychrometers to estimate both temperature and humidity conditions within hibernacula (Supplemental Figure 9). Humidity was measured as vapor pressure deficit (VPD), which measures the difference between the amount of moisture in the air and how much moisture the air could hold at the same temperature when saturated. Higher VPD values correspond to drier conditions.

For each hibernaculum, we calculated the mean early hibernation (November and December) temperature and VPD across all available years of data. We quantified the environmental conditions present early in hibernation because they are most associated with initial infection severity and ultimately survival probability (Hopkins et al., 2021; Langwig et al., 2016). Mean VPD values were skewed towards zero with a long right tail, so we log_10_-transformed the mean values before further analysis. To include untransformed VPD values of zero, a constant of 1×10^−5^ (same order of magnitude as the next-lowest non-zero VPD value) was added to all VPD values before transformation. Mean early hibernation temperature values were available for all 48 hibernacula and mean early hibernation VPD values were available for 34.

### Statistical analyses

#### Effect of population size on declines

To descriptively illustrate how colony classification captured differences in λ_i_ across epidemic phases, colony status, and their interaction, we constructed a generalized linear mixed model with gamma error distribution, log link function, and colony ID as a random effect (Model 1; all models are described in Supplemental Table 10). We explored associations between λ_i_ (response variable) and log_10_ colony size (N_i-T_ in equation 1) interacting with epidemic phase (predictors) using a generalized linear mixed model with gamma error distribution and a log link with colony ID as a random effect (Model 2). Colony size was confounded with colony status and was therefore used in a different model exploring λ_i_. We additionally assessed whether colony status was related to the size of the colony prior to the arrival of WNS by using a negative binomial model with log link function, the pre-WNS detection census value for each colony as the response variable, and colony status as the explanatory variable (Model 3).

#### Effects of environmental conditions on declines

To explore associations between environmental conditions within hibernacula and the response of Indiana bat colonies to WNS, we used a series of linear and generalized linear models. First, to test for the effect of mean early hibernation temperature, epidemic phase, and their interaction on the average annual colony growth value of a colony in an epidemic phase, λ_m_, we used a generalized linear model with a gamma error distribution and log link function (Model 4). The same model structure was used for a separate model to determine whether mean early hibernation VPD, epidemic phase, and their interaction influenced colony growth, λ_m_ (Model 5). Because environmental conditions may have influenced colony size prior to WNS arrival, we also examined the relationship between environmental conditions of hibernacula and pre-WNS detection colony size by using two negative binomial generalized linear models with log link functions and mean early hibernation temperature (Model 6) or VPD (Model 7) as the predictor and the latest pre-WNS colony size for each colony as the response variable. All linear models satisfied assumptions of Gaussian error distributions and homoscedastic error. All analyses were performed in R packages *stats* (R Core Team 2022), *glmmTMB* (Magnusson et al., 2017), and *MASS* (Ripley et al., 2013) in R v4.2.1.

#### Effect of little brown bat abundance on Indiana bat declines

Little brown bats (*Myotis lucifugus*) are a relatively abundant species in WNS-impacted communities and may contribute to community-wide disease dynamics (Laggan et al., 2023). However, because colony declines in little brown bats are also associated with the temperature and humidity of hibernacula, it is possible that any apparent associations between environmental conditions and Indiana bat declines could arise instead from the contribution of little brown bat abundance to Indiana bat survival and colony response. To identify a potential effect of little brown bat abundance on Indiana bat colony declines, we constructed a generalized linear mixed model with gamma error distribution and log link function with λ_i_ of Indiana bat colonies as the response variable, the log_10_-transformed colony size of little brown bats (N_i-T_ of little brown bats) interacting with epidemic phase as the explanatory variables, and colony ID as a random effect (Model 8). Thirty Indiana bat hibernacula had sufficient little brown bat census data to be included in Model 8.

#### Colony extirpation simulation

We performed a simulation to predict the proportion of the 231 colonies with known population status that would be extirpated by year 2030. We first classified each colony as “small” (≤ 66 individuals) or “large” (>66 individuals) based on its most recent colony census value prior to the detection of WNS within its hibernaculum, using the median colony size of 66 as the cut-off. If a pre-WNS census value was not available due to survey gaps, then we used the earliest census value post-WNS detection instead to classify the size of the colony. We constructed cumulative extirpation curves for each of these two datasets using colonies that were WNS-positive for at least seven years because the probability of becoming extirpated appeared to plateau following epidemic year 6. We fit logistic models to the two cumulative extirpation curves (Supplemental Figure 1, Supplemental Table 3) using the *stats* package in R (R Core Team 2022) and estimated the probability a colony was extirpated in epidemic year *t* if extant in epidemic year *t-1* as the difference between the epidemic year model estimates for both small and large colonies. We then simulated the annual probability of extirpation by 2030 (binomial variable, 1 = extirpated 0 = extant) using the most recent known status of each colony as well as the epidemic year of its most recent census. To simulate colony extirpation, each small or large colony that was extant in year *t-1* was assigned a new extirpation status in year *t* by drawing from a binomial probability distribution with the probability of extirpation corresponding to that calculated for each epidemic year using the cumulative extirpation curves for small and large colonies. The simulation was run for 1,000 iterations and summarized.

## Results

### General population trends

Population trends of Indiana bat colonies varied over time since pathogen arrival to their hibernacula (Figure 1, Supplemental Figure 2). Prior to the detection of WNS, colonies of Indiana bats were growing at an average rate of 0.9% annually. Following WNS detection within hibernacula, colonies declined significantly, with the largest declines in epidemic year two. Of the 231 colonies in our dataset, 56 (24.24%) were extirpated following the detection of WNS within their hibernacula and 53.57% of these extirpations occurred between epidemic years zero and two (Supplemental Figure 3). Additionally, 72 of the 231 colonies (31.16%) continued to decline following WNS arrival and never achieved an annual lambda, λ_i_, greater than or equal to 1.0 (stability) following WNS detection. The remaining 103 colonies (44.58%) were stable or growing for at least one year following the second year of the epidemic, and 47.57% of these had their first stable year by epidemic year three (Supplemental Figure 3). Population trends during the epidemic phase varied between extirpated, declining, and persisting colonies, with lower annual declines during the epidemic phase in persisting colonies compared to declining (β = 0.218 ± 0.043 SE, p < 0.001) or extirpated (β = 0.405 ± 0.082 SE, p < 0.001; Figure 1, Supplemental Table 4). Additionally, we found that extirpated colonies were, on average, significantly smaller than persisting (β = −3.1678 ± 0.356 SE, p < 0.001) or declining (β = −2.517 ± 0.389 SE, p < 0.001) colonies (Supplemental Figure 4, Supplemental Table 5). Pre-WNS arrival colony size in declining colonies was significantly lower than those in persisting colonies, but this effect was small (β = −0.651 ± 0.326 SE, p = 0.046). We did not detect any statistically significant association between either counts of Indiana bats or little brown bats at the same site on annual Indiana bat colony growth, λ_i_, regardless of epidemic phase (Supplemental Figure 5, 6, Supplemental Table 6, 7).

**Figure 1:**
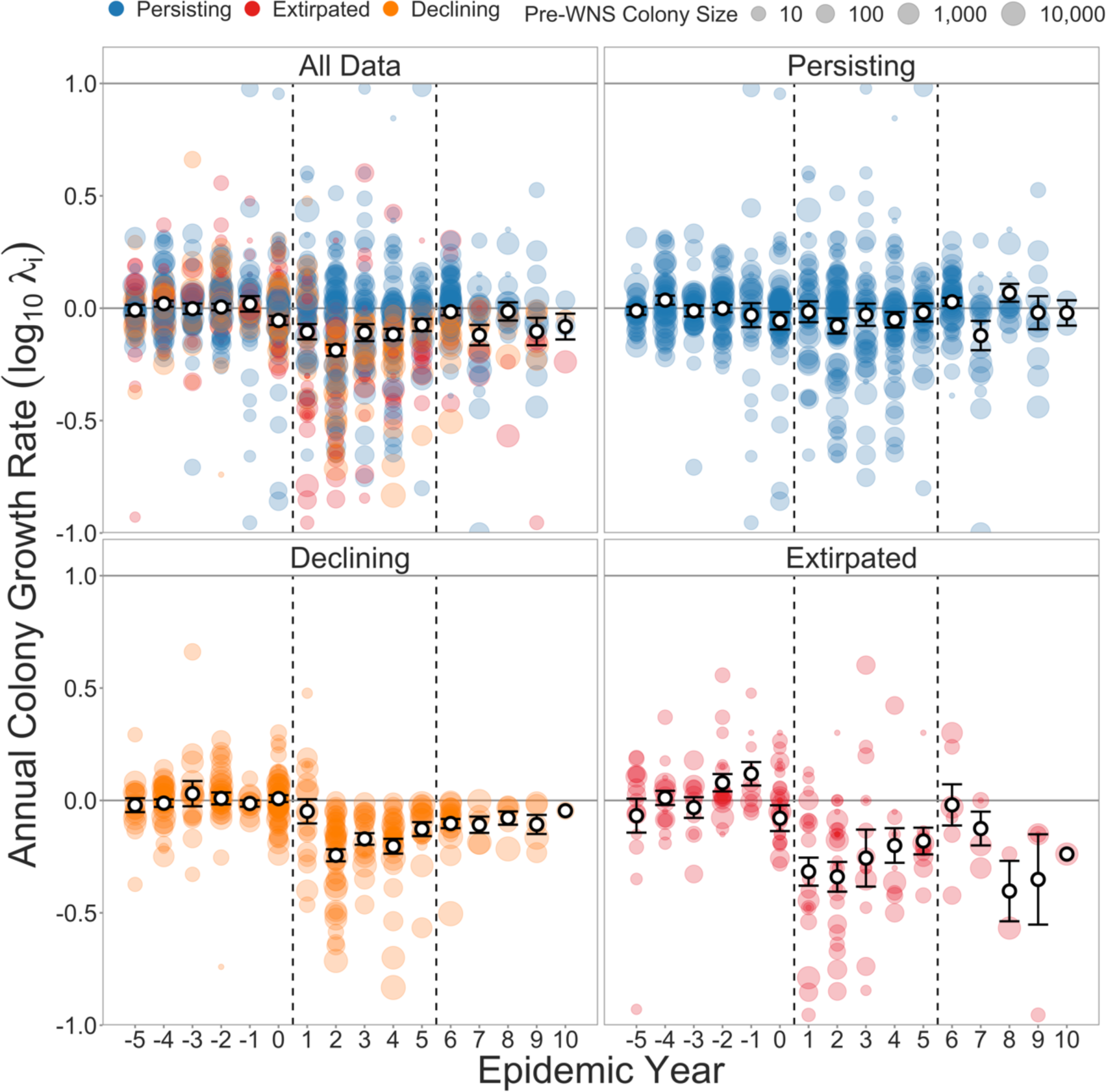
Population trends of Indiana bat colonies over time since detection of WNS. Epidemic year 0 on the x-axis corresponds to the year in which WNS was detected within a hibernaculum. Vertical dashed lines separate epidemic phases (left to right: pre-invasion, epidemic, established). Annual population growth rates of 0 on the y-axis correspond to population stability, and values below or above 0 indicate declining or growing colonies, respectively. Points are colored by colony status and size corresponds to pre-WNS arrival colony size. White points and error bars show mean values and one standard error.

### Colony extirpation simulation

Our simulation of future Indiana bat colony extirpations suggests that the rate of colony extirpation will slow in coming years (Figure 2). Most extirpations occur early in the epidemic and the probability of colony extirpation declines in later epidemic years (Supplemental Figure 1, 3). Therefore, our simulation predicts a decay in the rate of new extirpations, with 26.41% of total colonies (simulated range: 24.24 – 30.30%), or 61 of the 231 colonies in the dataset (simulated range: 56 – 70), extirpated by 2023. Because most extirpations occur early in the epidemic and most of the Indiana bat range has been affected by WNS for at least 5 years, this constitutes only an additional five extirpations from those recorded as of 2019. However, the number of predicted extirpations varied between small and large colonies. One average, 43.10% (range: 39.66 – 49.1%) of small colonies will be extirpated by 2030, corresponding to approximately 50 of the 116 small colonies in the dataset (range: 46 – 57), an increase of four extirpations from 2019. Conversely, only 9.57% (range: 8.70 – 12.17%) of large colonies will be extirpated, or approximately 11 of the 115 large colonies in the dataset (range: 10 – 14), an addition of a single extirpation from 2019.

**Figure 2:**
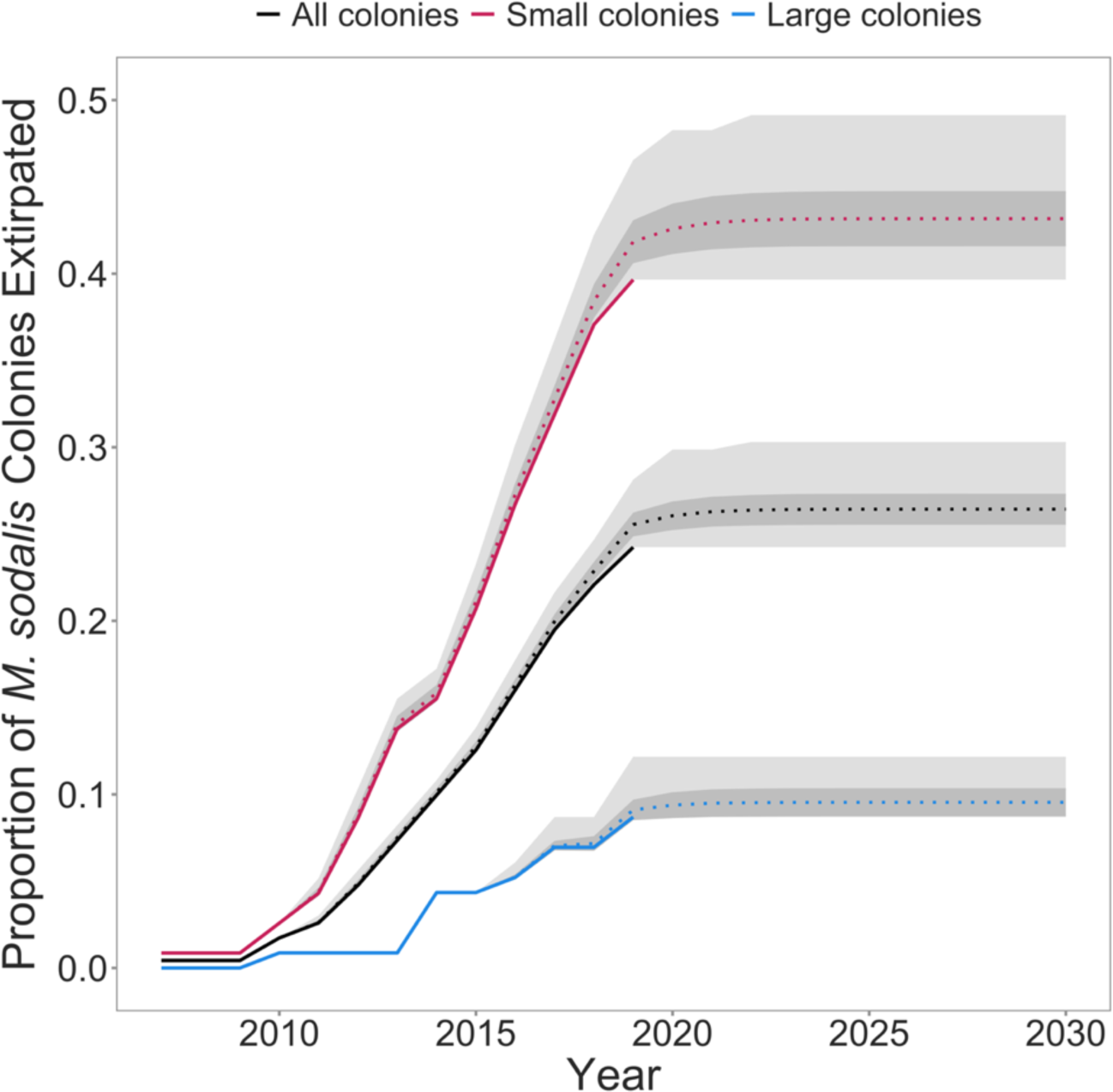
Simulated and empirical accumulation of extirpated Indiana bat colonies over time as a proportion of small, large, or all colonies. Solid lines show observed data and dotted lines show average simulated values for each year. The color of lines corresponds to dataset (black = all colonies, red = small colonies (≤ 66 individuals), blue = large colonies (> 66 individuals)). The dark gray shaded region corresponds to one standard deviation from the mean simulated value at each year. The light gray shaded region corresponds to the range of simulated values at each year.

### Environmental associations

Associations between early winter temperature conditions within hibernacula and population growth rates varied with epidemic phase (Figure 3A, Supplemental Table 8). We did not detect a statistically clear association between average early hibernation temperature and annual population growth of colonies prior to WNS arrival within hibernacula (temperature coefficient during Pre-invasion phase β = −0.012 ± 0.021 SE, p = 0.555). However, following the detection of WNS (Epidemic phase), we found that colonies roosting at colder early hibernation temperatures experienced more severe declines (temperature coefficient during Epidemic phase: β = 0.066 ± 0.023 SE, p = 0.005), a significantly different association than that observed in the Pre-invasion phase (interaction coefficient: β = −0.078 ± 0.031 SE, p = 0.013). Correspondingly, the estimated mean early hibernation temperature of extirpated hibernacula was 5.338 °C (± 0.708 SE, p = <0.001), colder than both persisting and declining colonies with estimates of 7.971 °C (± 0.322 SE, p = <0.001) and 8.022 °C (± 0.481 SE, p = <0.001), respectively (Figure 4A). Early winter temperature conditions of hibernacula used by persisting colonies did not significantly differ from those used by declining colonies (β = −0.051 ± 0.578 SE, p = 0.930). Following the establishment of *P. destructans* within hibernacula, the positive association between average early hibernation temperature and population growth rates weakened towards the neutral association observed prior to the WNS epidemic (temperature coefficient during Established phase: β = 0.037 ± 0.025 SE, p = 0.138), such that the association was not statistically different than pre-invasion (β = −0.050 ± 0.033 SE, p = 0.130) or epidemic phases (β = 0.028 ± 0.034 SE, p = 0.403). We found no statistically significant association between early hibernation temperature and pre-WNS Indiana bat colony sizes (β = −0.082 ± 0.054 SE, p = 0.126; Supplemental Figure 7).

**Figure 3:**
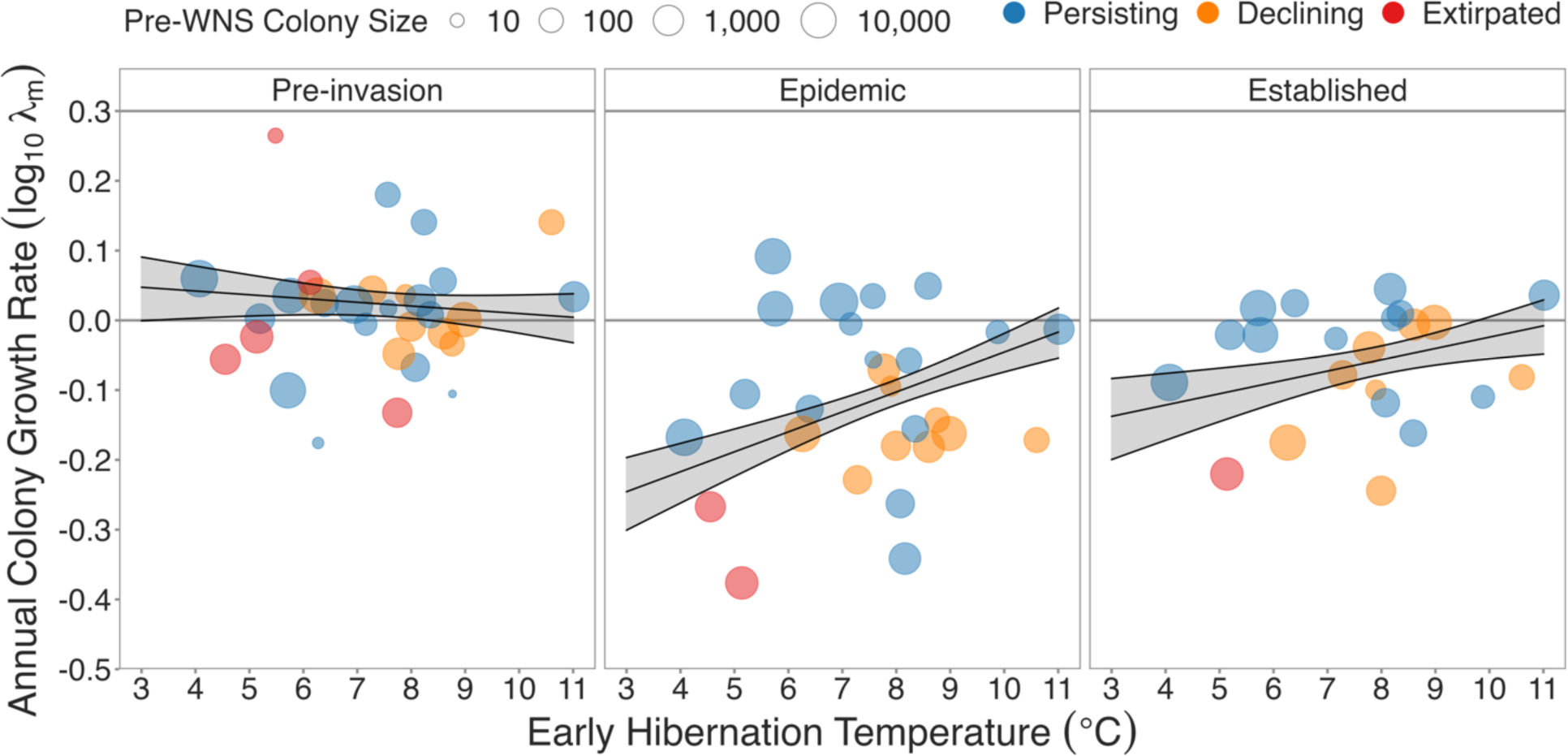
Annual colony growth rate over mean early hibernation temperature. Data is broken into separate panels by epidemic phase. A colony growth rate value of 0 corresponds to population stability, with values below or above 0 corresponding to colony decline or growth, respectively. Points are colored by colony status and point size corresponds to the most recent pre-WNS colony size. Solid black lines and shaded regions correspond to model estimates +/− one standard error of the model predictions.

**Figure 4:**
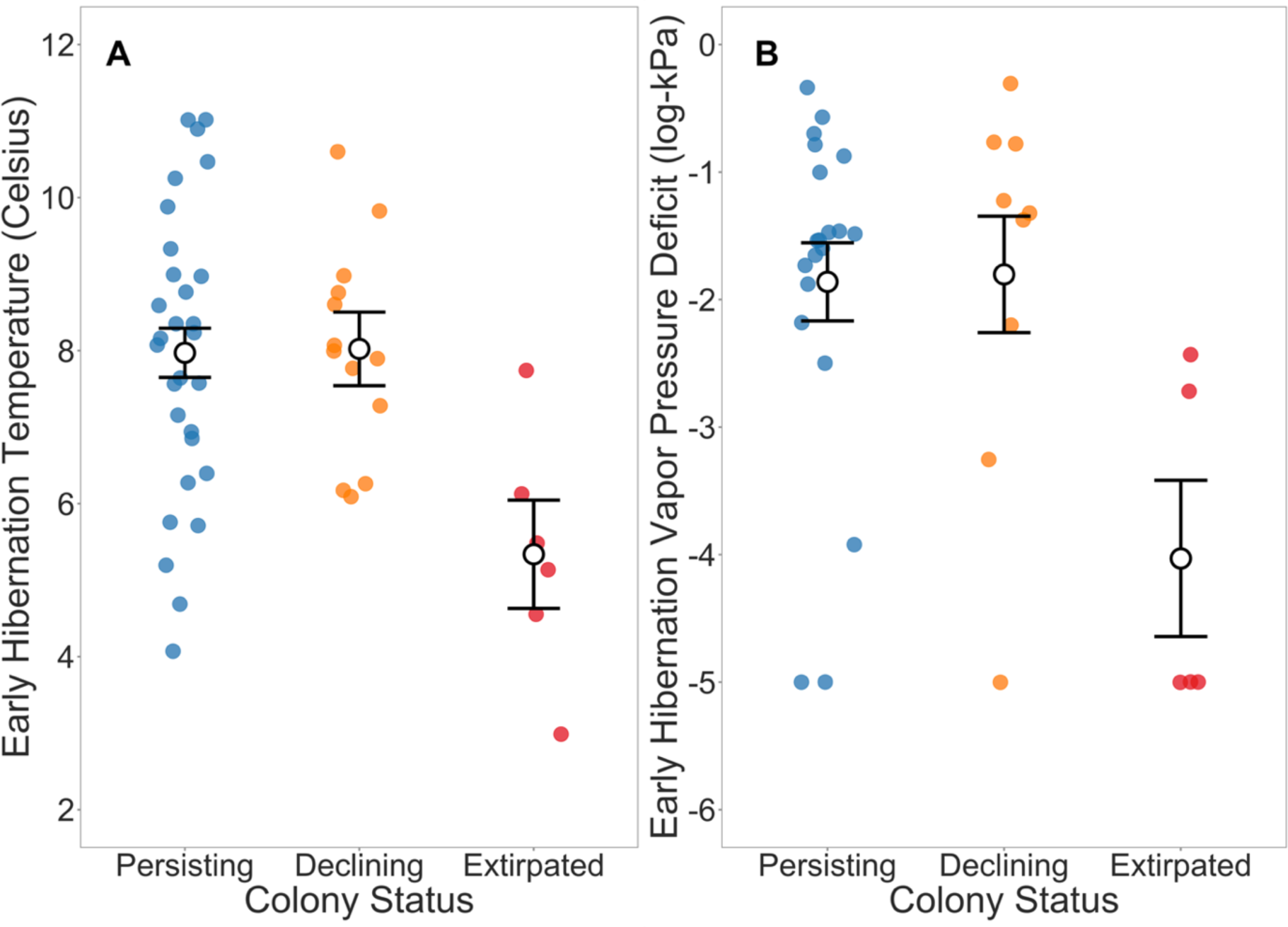
Effect of temperature and VPD on colony status. **A)** Mean early hibernation temperature and **B)** log_10_ vapor pressure deficit across persisting, declining, and extirpated colonies. Higher values of vapor pressure deficit correspond to drier conditions. Points are jittered and colored by colony status. White points and error bars correspond to model estimates +/− one standard error of predicted mean.

For vapor pressure deficit (VPD), we found no statistically clear relationships with colony growth, regardless of epidemic phase (Supplemental Figure 8, Supplemental Table 9). However, we found that VPD varied with site status, as extirpated colonies occurred in significantly more humid hibernacula than declining (β = −2.227 ± 0.764 SE, p = 0.007) or persisting colonies (β = −2.169 ± 0.685 SE, p = 0.003; Figure 4B). We detected a slight negative association between early hibernation VPD and pre-WNS Indiana bat colony sizes indicating that, prior to the arrival of WNS, drier hibernacula were used by relatively smaller colonies on average compared to humid hibernacula (β = −0.210 ± 0.093 SE, p = 0.025; Supplemental Figure 7).

## Discussion

We found that the factors associated with Indiana bat population dynamics shifted over epidemic time. Prior to the arrival of WNS, we did not detect an effect of colony size, hibernaculum temperature, or humidity on colony growth. Following the invasion of *P. destructans*, however, responses of Indiana bat colonies varied from complete extirpation to persistence, with smaller colonies being more prone to extirpation. In addition, colonies in colder hibernacula experienced more severe declines than those in relatively warm hibernacula and were more likely to be extirpated, a novel association that arose during the WNS epidemic. Colonies in relatively humid hibernacula were also at elevated risk of extirpation, potentially another association that arose following disease emergence. Once the pathogen became established in colonies, however, we found that the effects of environmental conditions weakened slightly, suggesting a possible return to the more neutral association observed prior to WNS arrival. Together, these data illustrate the potential of novel population stressors to shift the biotic and abiotic conditions optimal for population growth, an important consideration for the mitigation of their impacts.

Although population responses of Indiana bats were highly variable, many colonies in our dataset eventually stabilized in the presence of WNS. Colony persistence has been observed in other species of hibernating bats impacted by WNS in North America, such as the little brown bat (Dobony et al., 2011; Frick et al., 2015; Hoyt et al., 2021; Langwig et al., 2017, 2012), potentially the result of an interaction between evolutionary forces and favorable environmental conditions that allow for relatively high survival (Grimaudo et al., 2022; Hopkins et al., 2021; Langwig et al., 2017). Persisting Indiana bat colonies had less severe declines during the initial WNS epidemic than declining or extirpated colonies, potentially due to their larger colony sizes and warmer hibernaculum environments which buffered against unsustainable rates of mortality. However, stabilized colonies are not necessarily completely protected from extirpation, as they may be declining on average despite having a single year of colony stability or growth.

In accordance with prevailing theory, we found that extirpation risk was elevated in small colonies, potentially due to susceptibility to stochastic fadeout (Lande, 1993; Lande et al., 2003; Melbourne and Hastings, 2008) or demographic Allee effects (Gascoigne et al., 2009; Kramer et al., 2009; Stephens et al., 1999). Indiana bats are a highly gregarious species, historically having formed large clusters during hibernation (Clawson et al., 1980; Hardin and Hassell, 1970; Thomson, 1982), a behavior potentially beneficial for limiting energy expenditure during arousals from torpor (Boyles et al., 2008). Sufficiently low colony sizes may prevent conspecifics from forming clusters, increasing per-arousal energy expenditure already exacerbated by infection with *P. destructans*, causing down-stream effects on survival and ultimately leading to colony decline and extirpation. Further, the inability to form sufficiently large clusters may elevate emigration from reduced colonies, additionally contributing to their decline. If immigration to colonies occurs through individuals being led to hibernacula by conspecifics, then reduced colony sizes will further receive little immigration input, potentially slowing population growth in small colonies. Together, the consequences of reduced colony size may operate synergistically to elevate the risk of complete colony extirpation. More generally, our results suggest that pre-stressor population size, often a factor used to classify population risk, was positively associated with population resilience following pathogen arrival despite the potential for greater host density to amplify pathogen transmission.

Although small colonies were more likely to go extinct, we find that most colonies that will become extirpated have likely already done so due to the spread of WNS across the Indiana bat range. As the *P. destructans* invasion front advanced from its origin near Albany, New York, pathogen arrival was detected in Indiana bat colonies in every year between 2007 and 2017 in our dataset. However, all extant colonies are currently in later stages of the epidemic, when mortality is reduced and extirpation is less likely, resulting in a steep decline in the extirpation rate as the most recently infected colonies are either lost or stabilize. In the near future, declining colonies that have not yet been extirpated will either entirely collapse or settle into a stable population trajectory, ultimately determining the overall impact of WNS on Indiana bat colonies across their range.

We found that environmental conditions of hibernacula influenced Indiana bat colony response to WNS, an association that varied over epidemic time. In the years of high mortality and colony decline immediately following *P. destructans* arrival (epidemic phase), colonies using warmer hibernacula experienced lower declines than those in colder hibernacula. The role of temperature appeared to have arisen due to the emergence of WNS, as we did not detect the association prior to detection of the disease within hibernacula, illustrating the ability of novel infectious diseases to alter associations between environmental conditions and population growth. The effect of temperature extended to extirpation risk, as extirpated colonies used significantly colder hibernacula than extant colonies. Furthermore, while we did not detect an association between VPD within hibernacula and colony growth rates, we did find that extirpated colonies typically used more humid hibernacula than extant colonies, suggesting that humidity may additionally impact colony growth dynamics following WNS emergence. Importantly, we detected no association between pre-WNS colony size and temperature conditions within hibernacula, and a weak negative one for VPD, indicating that environmental conditions are unconfounded with colony size and operate independently to drive colony responses to WNS. Finally, while we did not investigate the role of temporal or spatial variation in temperature or humidity conditions within hibernacula on population response to WNS, it represents an additional potential driver of disease processes in this system. Future research is warranted to explore how variation in environmental conditions might influence disease processes from the individual- to population-level impacts on bat communities with WNS.

The positive association between hibernaculum temperature and colony response to WNS is opposite of that reported for the little brown bat, in which warmer hibernacula have higher disease severity and declines (Grieneisen et al., 2015; Hopkins et al., 2021; Langwig et al., 2016, 2012; Lilley et al., 2018), likely due to increased pathogen growth under those conditions (Grieneisen et al., 2015; Hopkins et al., 2021; Johnson et al., 2014; Langwig et al., 2016; Verant et al., 2012). We did not detect an association between little brown bat abundance within hibernacula and Indiana bat colony growth, suggesting that Indiana bat population declines at colder temperatures are not due to larger little brown bat colonies in cold hibernacula increasing environmental pathogen contamination (Laggan et al., 2023) or disturbance from torpor (Turner et al., 2014). Instead, this discrepancy might arise if the energy expended during arousals from torpor, which are more frequent in bats with WNS (Lilley et al., 2016; Reeder et al., 2012; Warnecke et al., 2012), is more important for surviving the disease for Indiana bats than little brown bats. The gregarious nature of Indiana bats and their propensity to form large clusters, which may further reduce cost of arousals from torpor (Boyles et al., 2008), lend possible credence to this explanation. Energy expended during arousals from torpor accounts for the majority of that spent during hibernation and is reduced under warmer conditions (Boyles and Willis, 2010; Thomas et al., 1990). The reduced cost of arousals in warm hibernacula potentially allowed Indiana bats to conserve enough energy to survive WNS over winter, buffering their overall declines. In cold hibernacula, however, the cost of additional arousals due to WNS may have been too great and resulted in premature energy depletion and mortality. For little brown bats, the disruption of water balance may instead be the physiological pathway that contributes most to mortality (Ben-Hamo et al., 2013; Cryan et al., 2010; Grimaudo et al., 2022; Thomas and Cloutier, 1992; Webb et al., 1995), which hypothetically scales positively with pathogen load. Given that the growth of *P. destructans* increases with hibernaculum temperature (Marroquin et al., 2017; Verant et al., 2012), little brown bat colonies in warm hibernacula could experience greater and unsustainable rates of water loss, resulting in their positive association between hibernaculum temperature and colony decline whereas the opposite exists for Indiana bats. Future research is warranted in identifying the physiological or ecological underpinnings of the opposite responses of Indiana and little brown bat colonies to hibernaculum temperatures when challenged by WNS.

The difference between Indiana and little brown bats in their colony response to WNS illustrates that in novel infectious disease systems with multiple host species, there may be species-specific differences in how environmental conditions influence population response to pathogen invasion. To mitigate the impact of invading pathogens on host populations, management strategies that consider inherent differences among host species are essential. For example, habitat manipulations meant to cool or warm hibernacula and benefit a target bat species could inadvertently harm another, so careful consideration of the bat communities present is essential (Boyles et al., 2023). Alternatively, the acquisition and conservation of diverse hibernacula critical to each species, such as those that offer protective microclimates or where colony populations are stabilizing, should be considered a high priority. Ultimately, the appropriate management strategy will be dependent on the community, environment, and disease conditions present, and management flexibility should reflect this potential for variation.

This study highlights how existing associations between biotic and abiotic conditions and population growth can be altered by novel stressors, including emerging infectious diseases. To protect vulnerable host populations from disease-driven extirpation, conservation measures should reflect these changing associations and consider differences among host species in shared habitat. Furthermore, it is crucial to conserve diverse habitats available to host species, as novel associations may arise and offer refugia from severe disease impacts. However, determining how to balance the conservation of habitat that is beneficial to one species while potentially detrimental to another (i.e. ecological traps), such as the case with thermal conditions of Indiana and little brown bat hibernacula, deserves additional research. As novel host—pathogen interactions continue to arise, the successful mitigation of their impacts will be dependent on our ability to characterize and respond to how they interact with the unique ecological context in which they arise.

## Acknowledgements

We thank those who contributed to the collection of census and environmental data across all participating agencies and organizations, without whom this study would not have been possible. Any use of trade, firm, or product names is for descriptive purposes only and does not imply the endorsement by the U.S. Government. The findings and conclusions in this article of those of the authors and do not necessarily represent the views of the U.S. Fish and Wildlife Service.

## Supplemental material

**Supplemental Figure 1:**
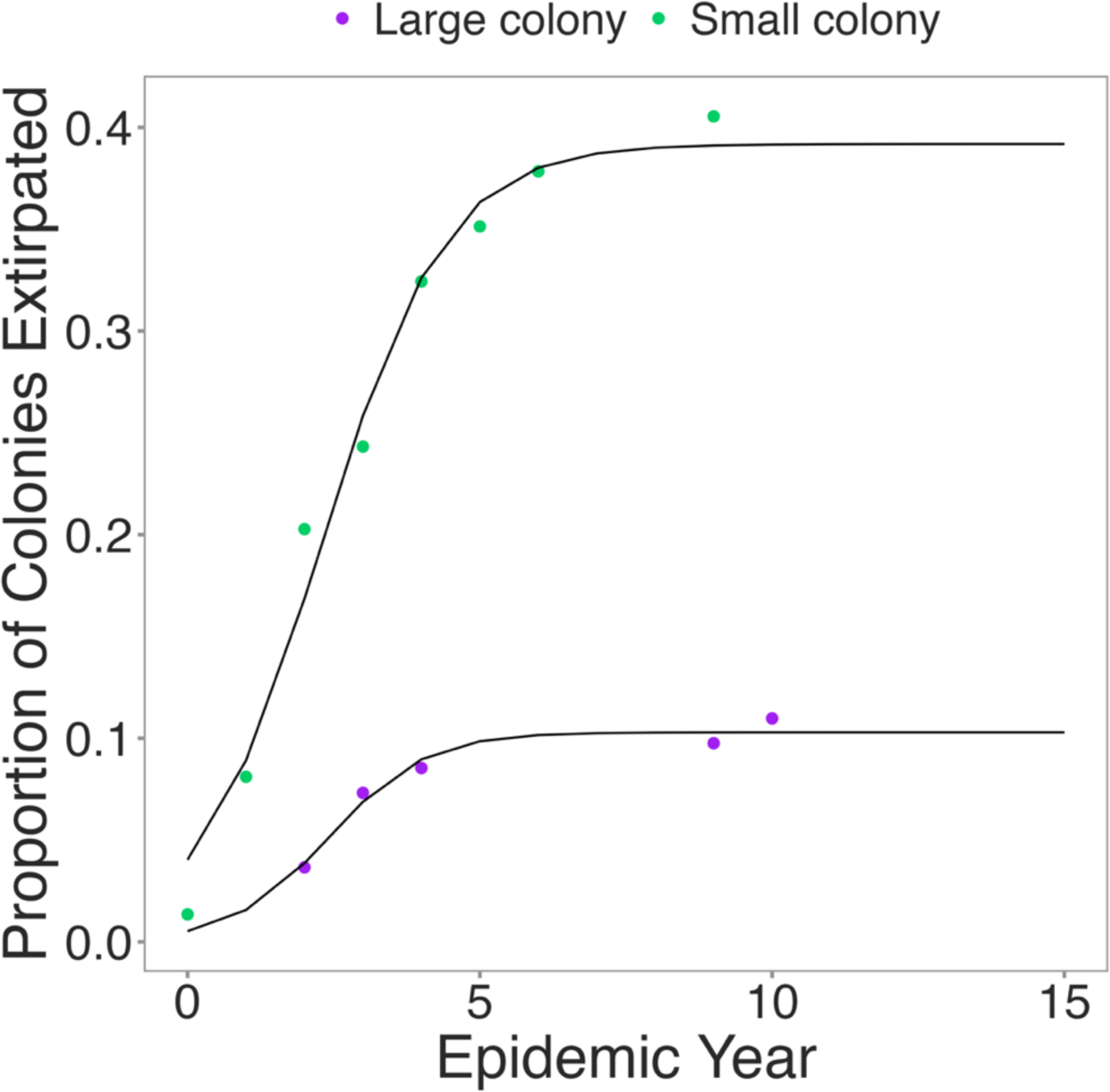
Plot of cumulative extirpations for both small (pre-WNS detection colony size of 66 or fewer individual *M. sodalis*) and large (pre-WNS detection colony size greater than 66 individual *M. sodalis*) colonies that have been WNS-positive for seven or more years (n = 74 small, 82 large colonies). Solid lines are estimates of logistic growth models fit to the cumulative extirpation data, from which the simulation extirpation probabilities were calculated.

**Supplemental Figure 2:**
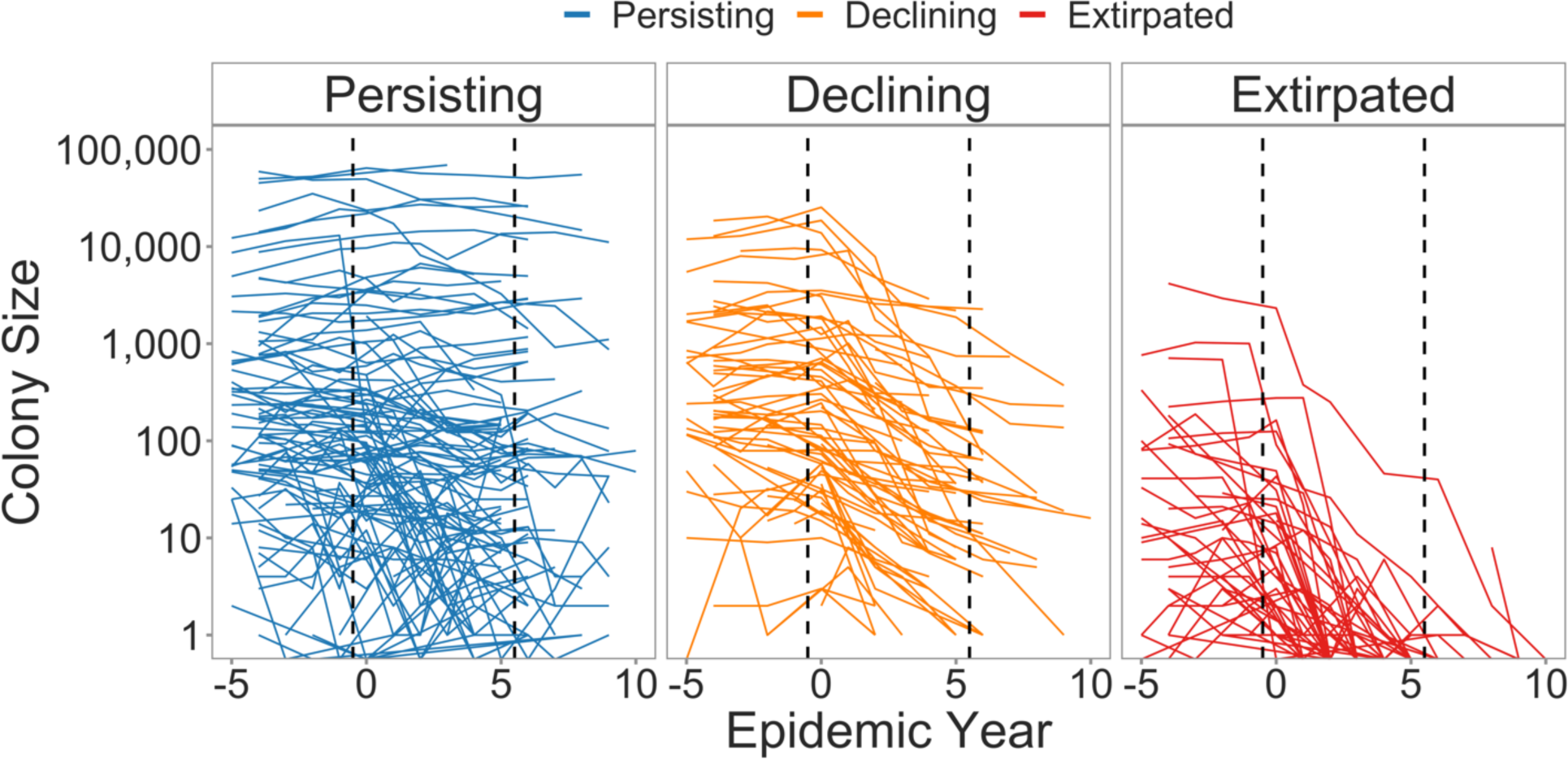
Colony sizes of persisting, declining, and extirpated colonies over time since WNS detection. Epidemic year 0 on the x-axis corresponds to the year in which WNS was detected. Vertical dashed lines separate epidemic phases (pre-invasion, epidemic, established).

**Supplemental Figure 3:**
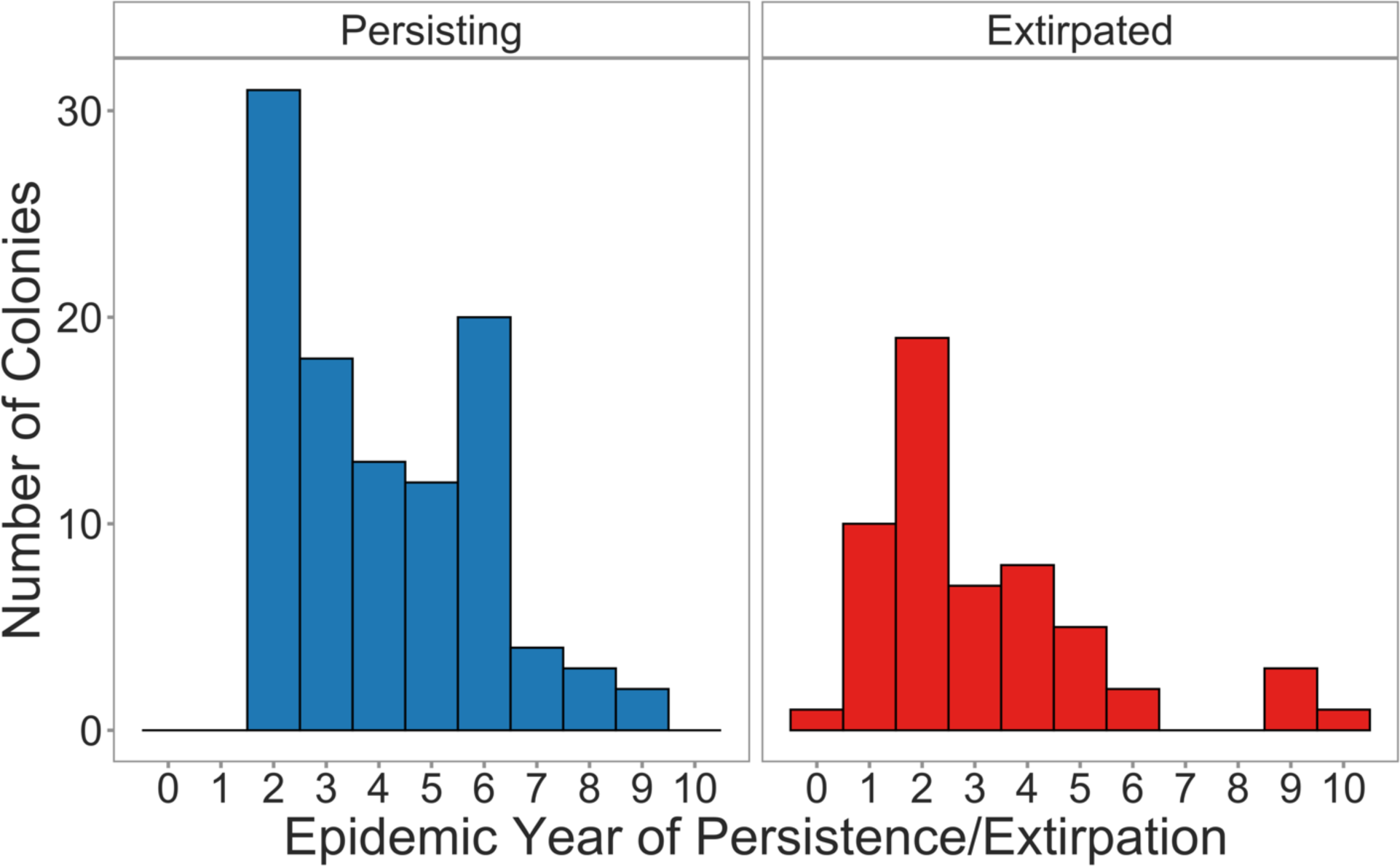
Histogram of epidemic year in which stability or extirpation first occurs in persisting and extirpated colonies, respectively. Years of stability in epidemic year 0 and 1 were not considered true stability events, as *P. destructans* is at sufficiently low prevalence at that point that the incidence of spurious population growth outside of the context of WNS is high.

**Supplemental Figure 4:**
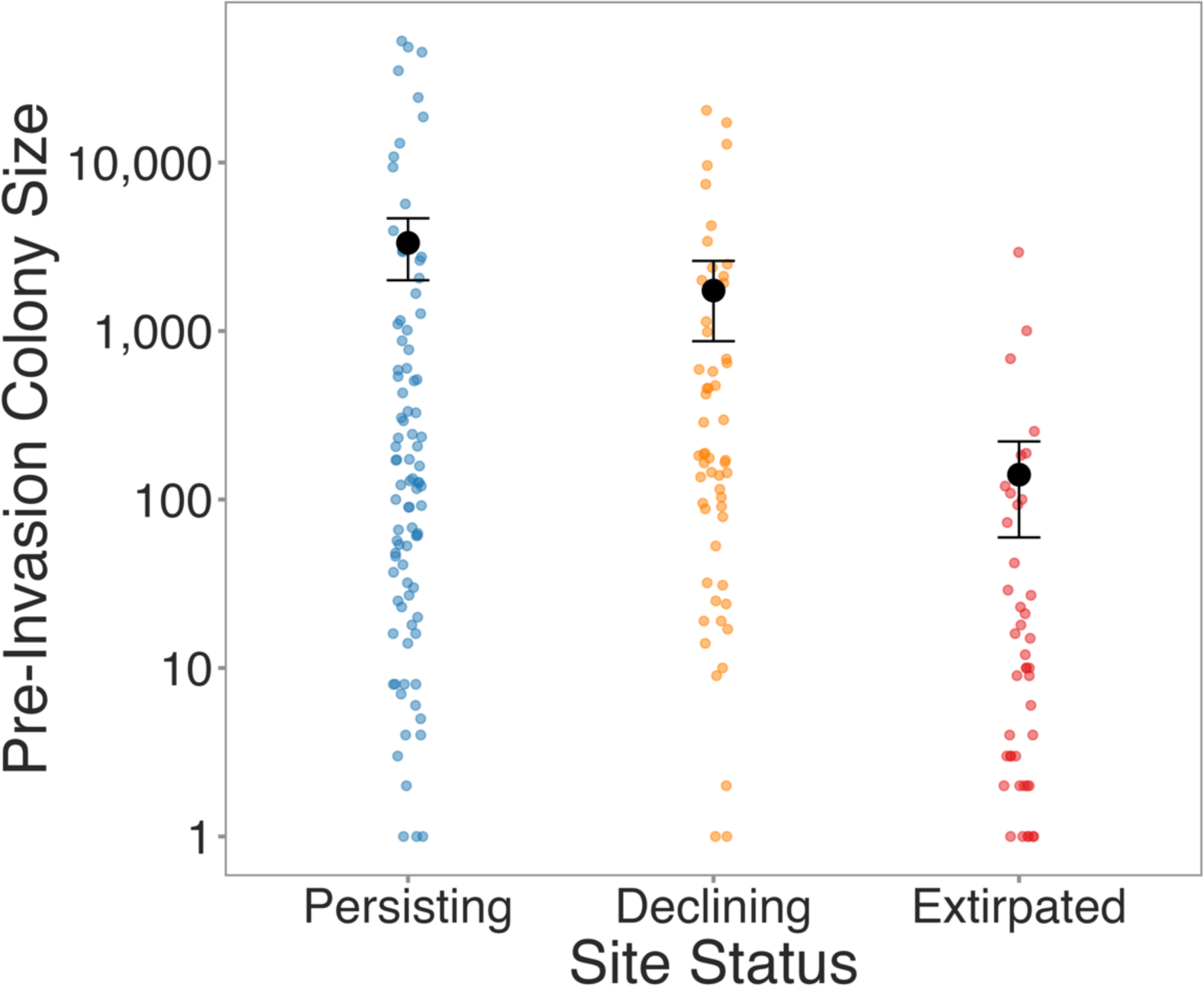
Most recent colony sizes of persisting, declining, and extirpated colonies prior to the detection of WNS. Points are jittered and colored by site status.

**Supplemental Figure 5:**
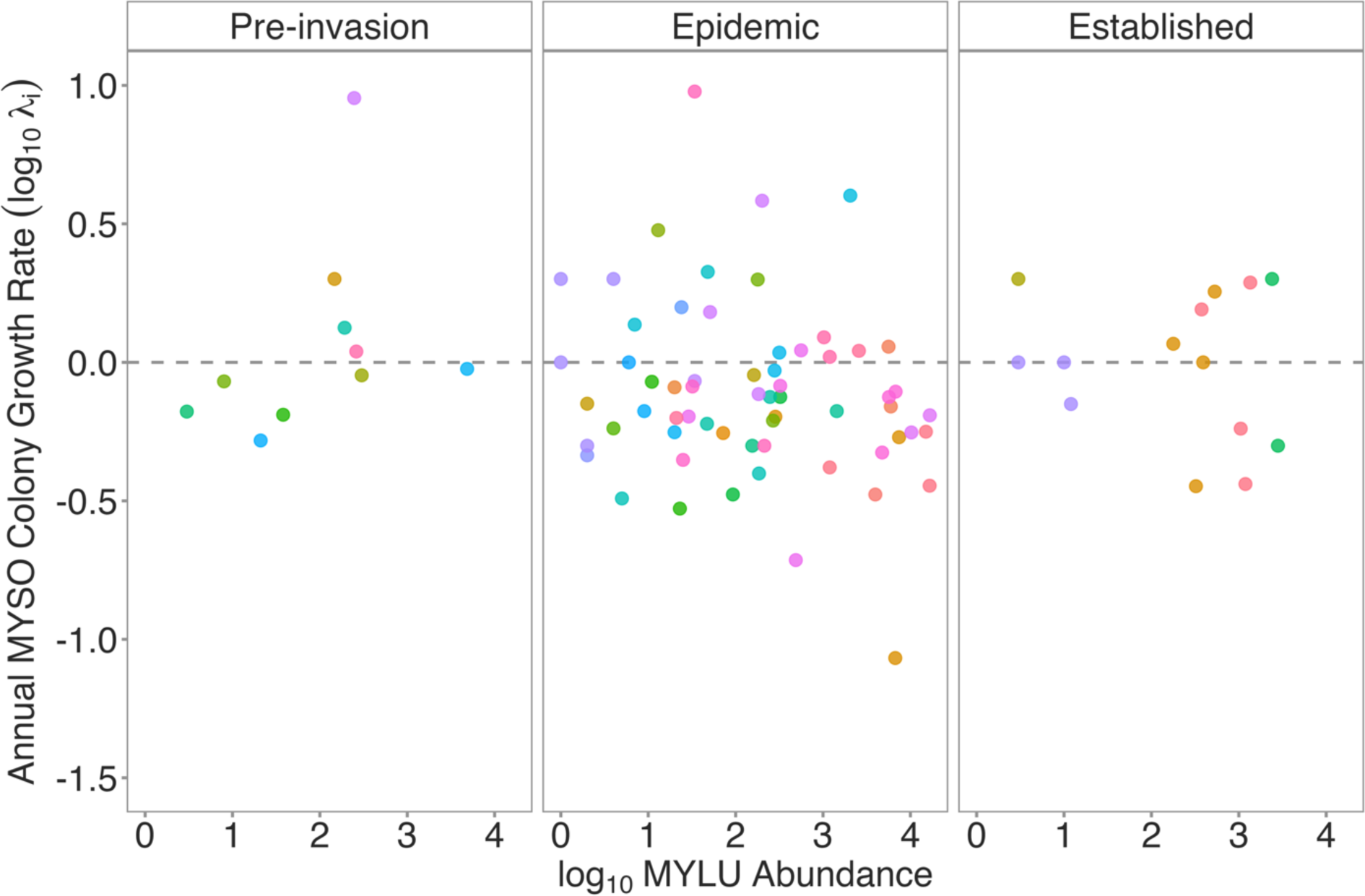
Annual Indiana bat (MYSO) colony growth rate by the log_10_ little brown bat (MYLU) abundance within the same hibernaculum in the previous winter. A colony growth value of 0 corresponds to colony stability, with values below or above 0 indicating colony decline and growth, respectively. Points are colored by unique hibernacula.

**Supplemental Figure 6:**
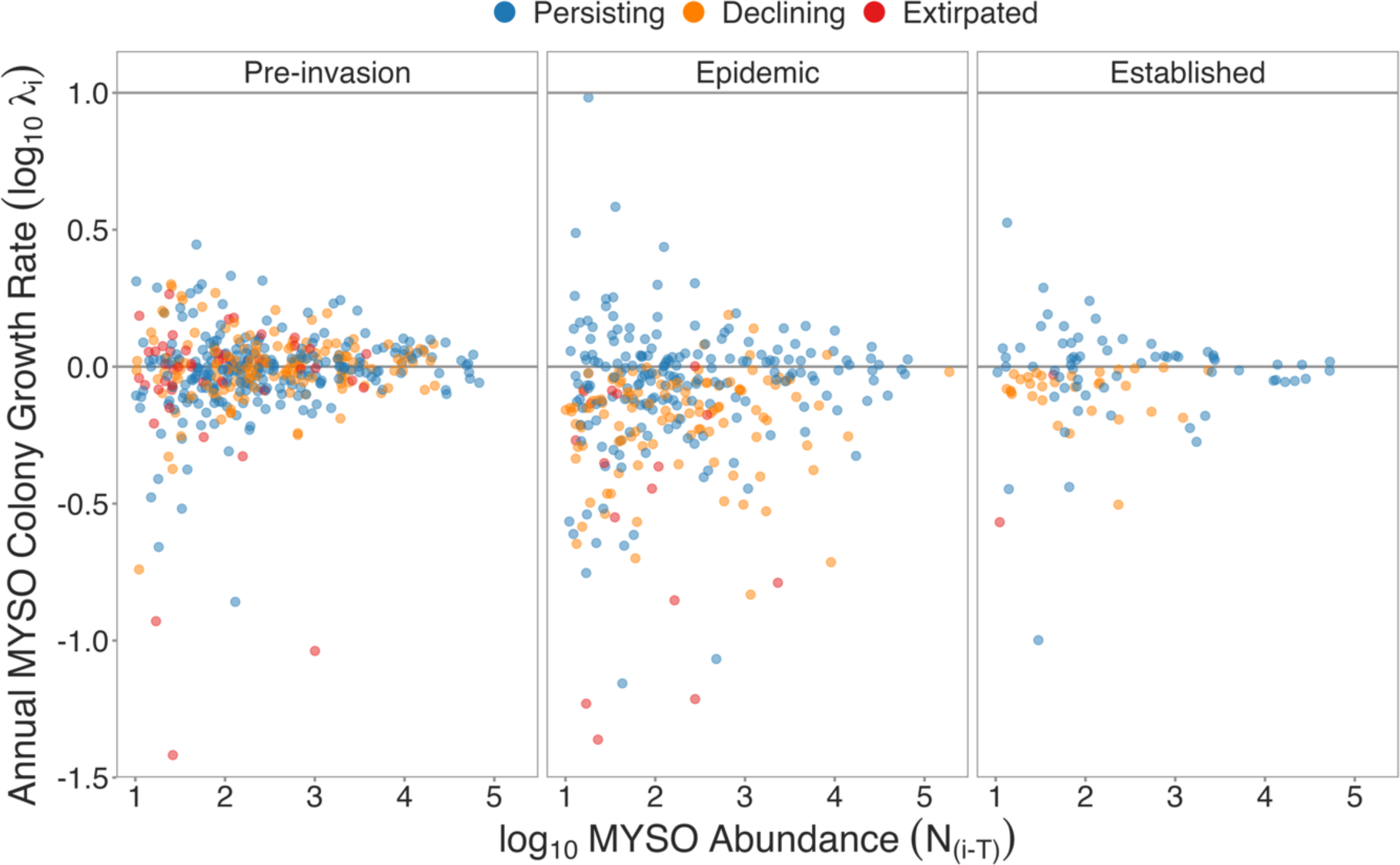
Annual Indiana bat (MYSO) colony growth rate by the log_10_ Indiana bat abundance within the same hibernaculum in the previous winter. A colony growth value of 0 corresponds to colony stability, with values below or above 0 indicating colony decline and growth, respectively. Points are colored by colony status.

**Supplemental Figure 7:**
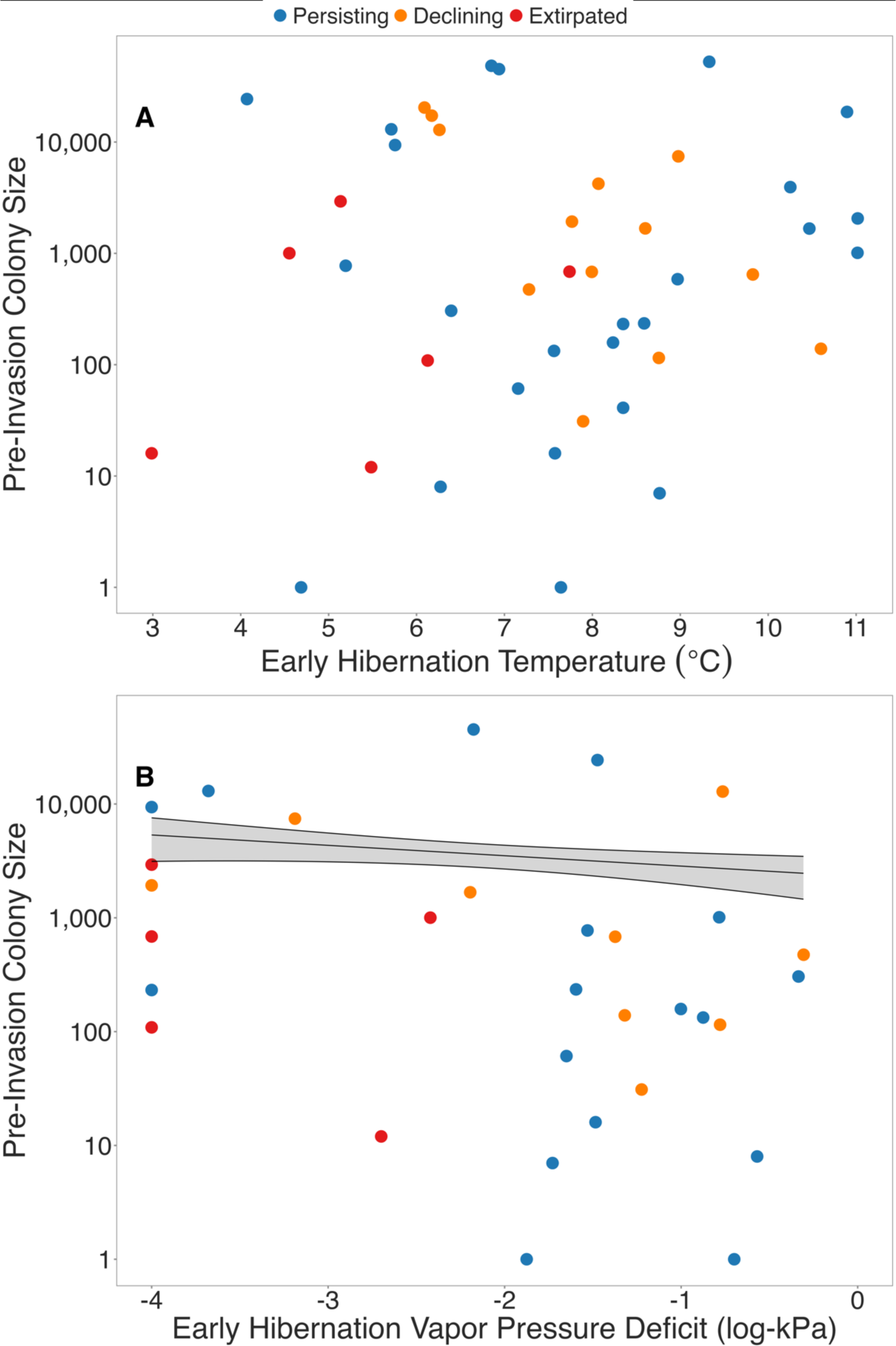
Association between pre-WNS Indiana bat colony sizes and **A)** mean early hibernation temperature (Celsius) and **B)** log_10_ vapor pressure deficit. Points are colored by colony status. Solid line and shaded region correspond to model estimates +/− one standard error of estimated mean.

**Supplemental Figure 8:**
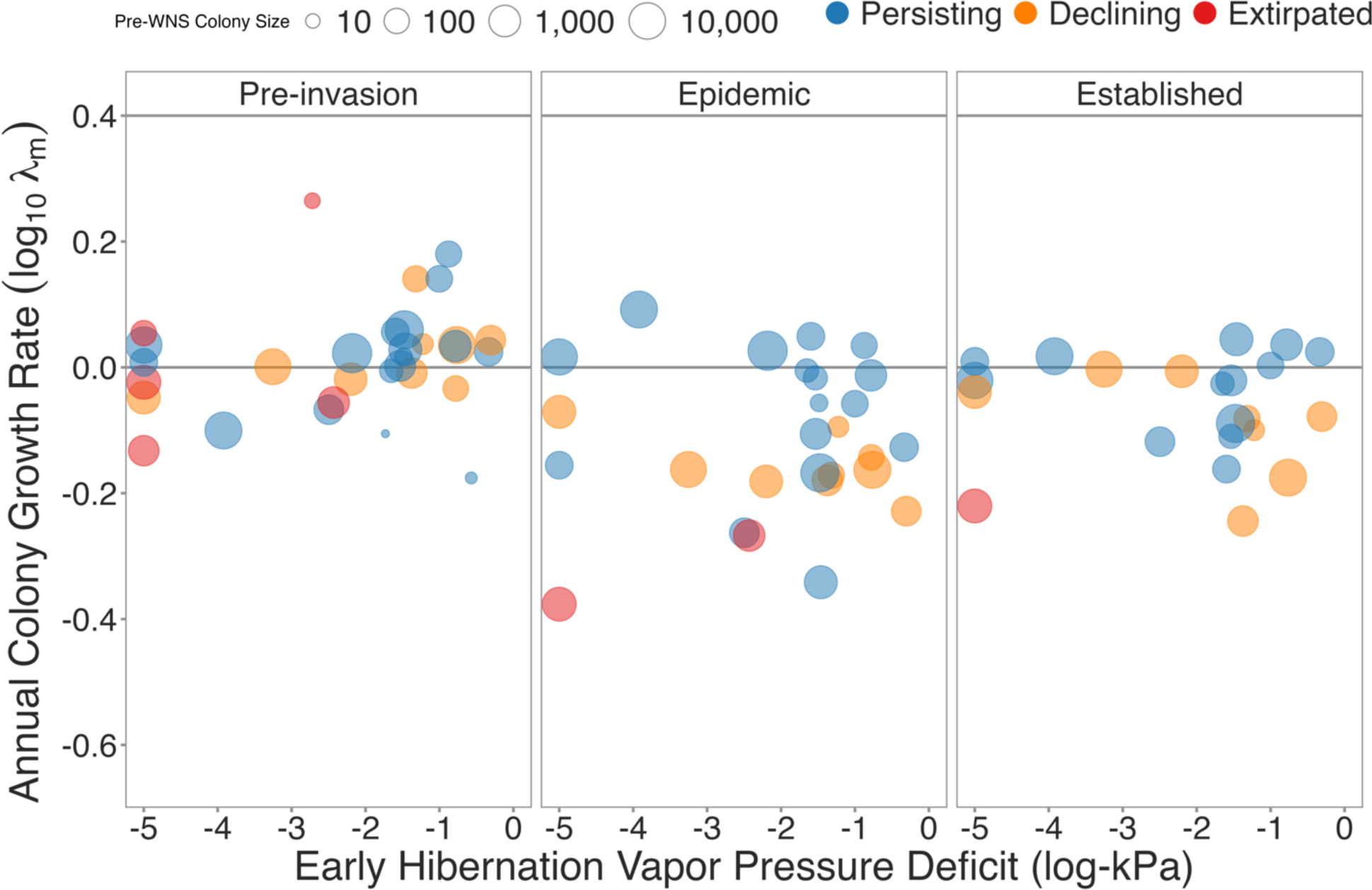
Annual colony growth rate over mean early hibernation vapor pressure deficit. Lower VPD values correspond to more humid conditions. Data is broken into separate panels by epidemic phase. A colony growth rate value of 0 corresponds to population stability, with values below or above 0 corresponding to colony decline or growth, respectively. Points are colored by colony status and point size corresponds to the most recent pre-WNS colony size.

**Supplemental Figure 9:**
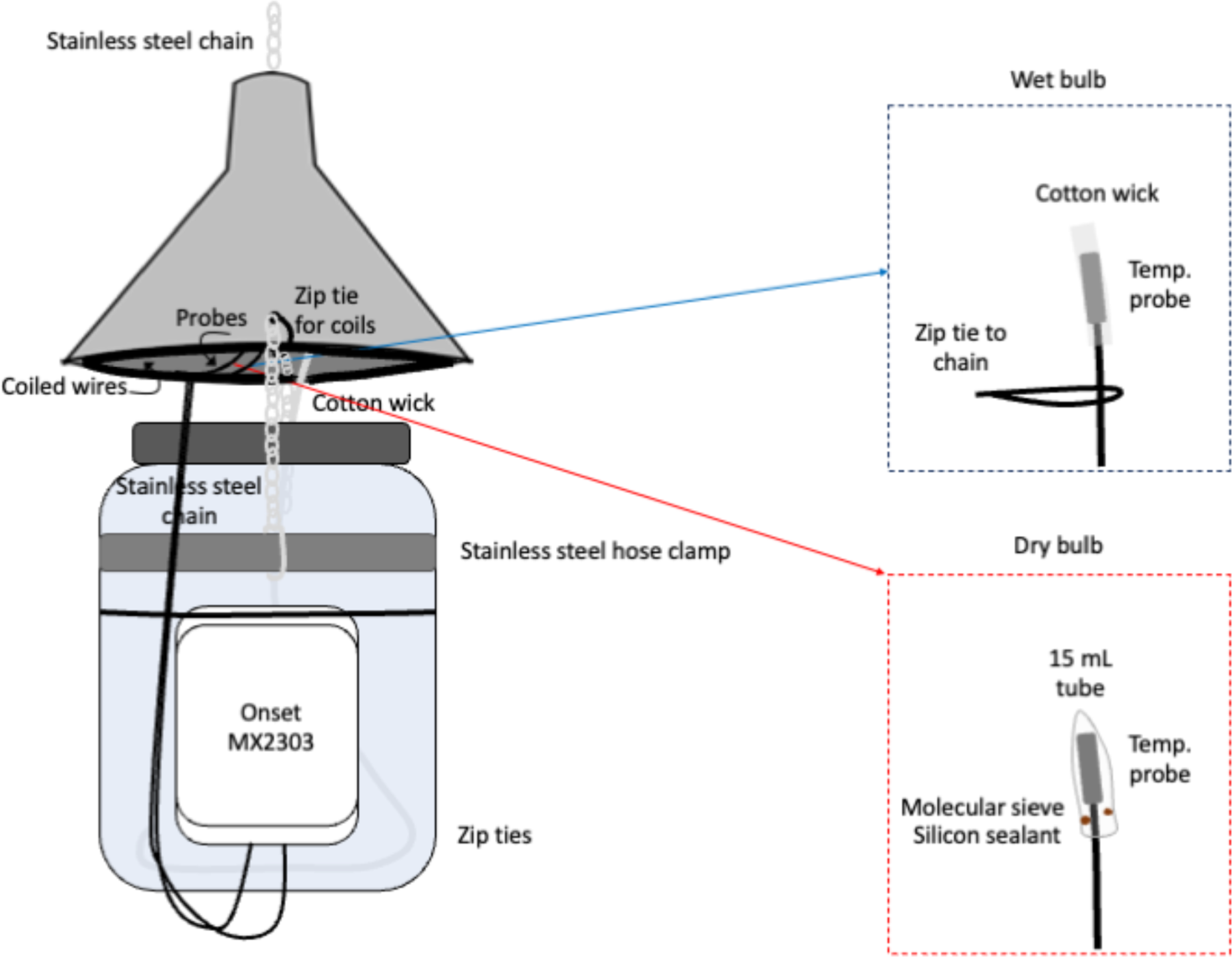
Schematic of wet-bulb psychrometer used in this study. Wet-bulb psychrometers compare the temperature of wet bulbs and dry bulbs to estimate the vapor pressure deficit (VPD), or the difference between the measured vapor pressure and the vapor pressure at saturation at the given temperature. High values of VPD indicate low vapor pressure relative to what could be achieved at saturation, indicating dry environments. Low values of VPD indicate near- or at-saturation vapor pressure, corresponding to humid conditions.

## Supplemental Tables

**Supplemental Table 1:** summary of origin of all Indiana bat count data we analyzed, including number of hibernacula per state, the range of years represented in their data, and the number of winter years of census data contributed to the dataset. The total winter years of count data contributed by each state was the number of census values from all hibernacula across all years of data. Also contains the number of hibernacula per state with both census and environmental logger data available, as well as the logger years of data from those hibernacula.

| State | Number of hibernacula with census data | Range of years of census data | Total winter years of census data (% of all census data) | Number of hibernacula with both census and logger data (logger years) |
| --- | --- | --- | --- | --- |
| Alabama | 8 | 2009—2019 | 39 (3.14%) | 0 (0) |
| Arkansas | 8 | 2009—2019 | 39 (3.14%) | 1 (0.17) |
| Illinois | 9 | 2009—2022 | 48 (3.87%) | 3 (13.99) |
| Indiana | 19 | 2007—2019 | 113 (9.11%) | 8 (10.43) |
| Kentucky | 75 | 2006—2019 | 345 (27.8%) | 4 (1.32) |
| Missouri | 36 | 2005—2019 | 137 (11.04%) | 3 (6.83) |
| North Carolina | 2 | 2008—2014 | 6 (0.48%) | 0 (0) |
| New Jersey | 2 | 2004—2018 | 13 (1.05%) | 0 (0) |
| New York | 16 | 2003—2019 | 106 (8.54%) | 8 (4.79) |
| Ohio | 2 | 2007—2016 | 11 (0.89%) | 0 (0) |
| Pennsylvania | 15 | 2004—2019 | 52 (4.19%) | 2 (0.33) |
| Tennessee | 28 | 2007—2019 | 137 (11.04%) | 4 (1.00) |
| Virginia | 15 | 2005—2019 | 76 (6.12%) | 4 (1.32) |
| Vermont | 2 | 2003—2017 | 6 (0.48%) | 1 (0.13) |
| West Virginia | 28 | 2005—2019 | 113 (9.11%) | 10 (10.42) |

**Supplemental Table 2:** summary of the amount of environmental data collected by each logger model in this study.

| Logger model | Total logger years of data (% of all environmental data) |
| --- | --- |
| Onset HOBO Pro | 4.21 (8.26%) |
| Onset HOBO Pro V2 | 28.86 (56.70%) |
| Onset HOBO MX2303 | 5.52 (10.84%) |
| Onset HOBO XT | 10.92 (21.45%) |
| Onset HOBO Pendant | 0.17 (<1%) |
| Maxim Integrate iButton | 0.37 (<1%) |
| Madgetech TransiTempII-RH | 0.33 (<1%) |
| Unknown | 0.51 (1%) |

**Supplemental Table 3:**
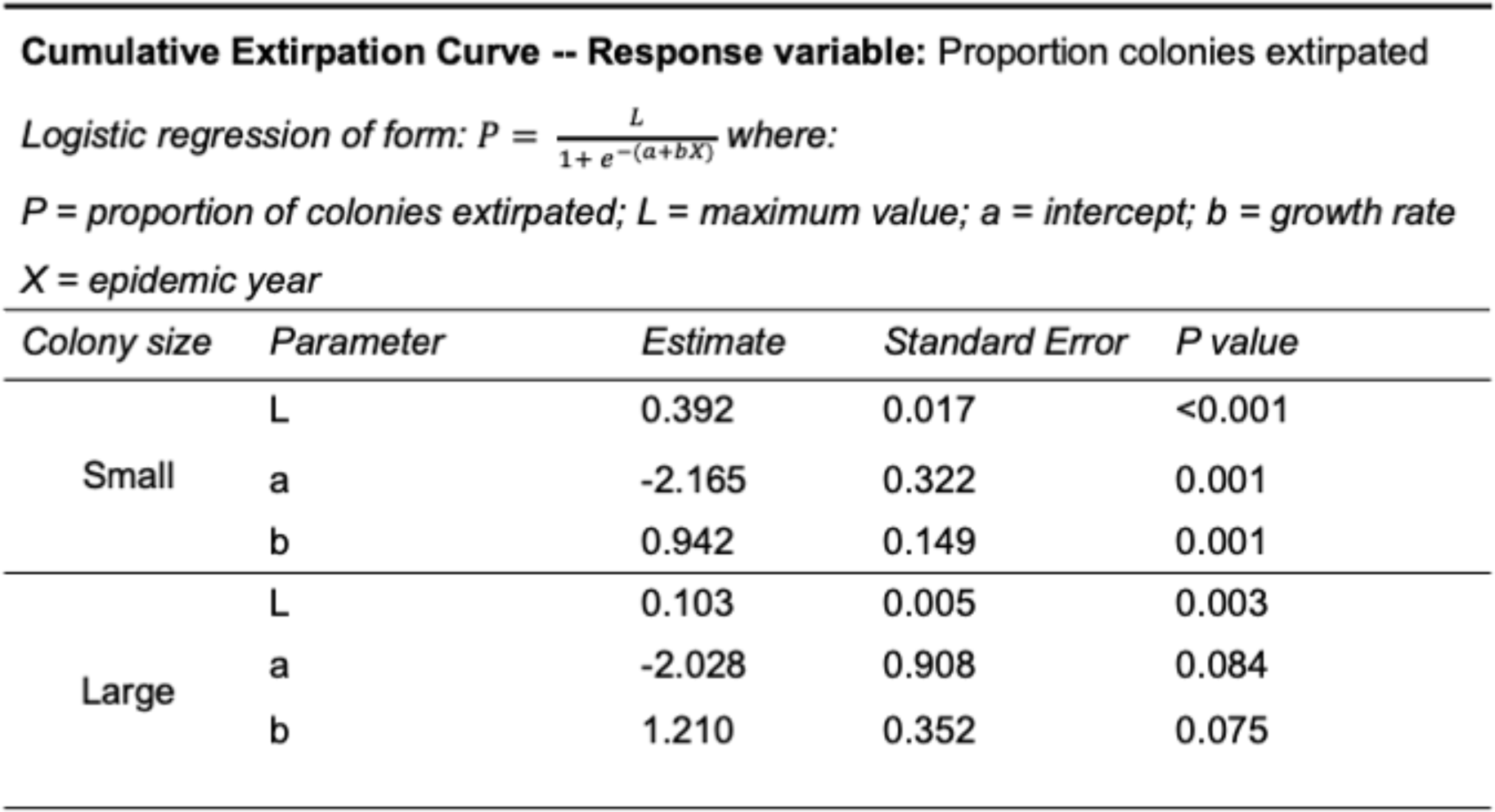
summary of the estimated parameters of a logistic regression fit to proportional cumulative colony extirpation data.

| Cumulative Extirpation Curve -- Response variable: Proportion colonies extirpated |  |  |  |  |
| --- | --- | --- | --- | --- |
| Logistic regression of form: $P = \frac{L}{1 + e^{-(a+bX)}}$ where: | | | | |
| *P* = proportion of colonies extirpated; *L* = maximum value; *a* = intercept; *b* = growth rate |  |  |  |  |
| *X* = epidemic year |  |  |  |  |
| Colony size | Parameter | Estimate | Standard Error | P value |
| Small | L | 0.392 | 0.017 | <0.001 |
|  | a | -2.165 | 0.322 | 0.001 |
|  | b | 0.942 | 0.149 | 0.001 |
| Large | L | 0.103 | 0.005 | 0.003 |
|  | a | -2.028 | 0.908 | 0.084 |
|  | b | 1.210 | 0.352 | 0.075 |

**Supplemental Table 4:** statistical output of a generalized linear mixed model exploring the association between epidemic phase, colony status, and their interaction with annual colony growth rate.

| **Model 1 -- Response variable: Annual colony growth rate ($\lambda_i$)** |  |  |  |
| --- | --- | --- | --- |
| *Generalized linear mixed model with gamma error distribution, log link function, random effect of colony ID (N = 857, groups = 182)* |  |  |  |
| *Model reference level: Epidemic phase(Pre-invasion), Colony status(Persisting)* |  |  |  |
| *Term* | *Estimate* | *Standard Error* | *P value* |
| Intercept | 0.028 | 0.032 | 0.386 |
| Epidemic phase(Epidemic) | -0.039 | 0.041 | 0.353 |
| Epidemic phase(Established) | 0.005 | 0.061 | 0.929 |
| Colony status(Extirpated) | -0.019 | 0.076 | 0.798 |
| Colony status(Declining) | 0.001 | 0.053 | 0.990 |
| Epidemic phase(Epidemic):Colony status(Extirpated) | -0.385 | 0.112 | <0.001 |
| Epidemic phase(Established):Colony status(Extirpated) | -0.521 | 0.314 | 0.097 |
| Epidemic phase(Epidemic):Colony status(Declining) | -0.219 | 0.068 | 0.001 |
| Epidemic phase(Established):Colony status(Declining) | -0.233 | 0.106 | 0.028 |

**Supplemental Table 5:** statistical output of a linear model exploring the association between colony status and pre-WNS arrival colony size.

| *Term* | *Estimate* | *Standard Error* | *P value* |
| --- | --- | --- | --- |
| Intercept | 8.112 | 0.204 | <0.001 |
| Colony status(Declining) | -0.651 | 0.326 | 0.046 |
| Colony status(Extirpated) | -3.168 | 0.358 | <0.001 |

**Supplemental Table 6:** statistical output of a generalized linear mixed model exploring the association between Indiana bat colony size in year *i*-1 with the annual colony growth to year *i*.

| *Term* | *Estimate* | *Standard Error* | *P value* |
| --- | --- | --- | --- |
| Intercept | -0.011 | 0.061 | 0.859 |
| $\log_{10}$ census value ($N_{i-1}$) | 0.012 | 0.024 | 0.612 |
| Epidemic phase(Epidemic) | -0.160 | 0.088 | 0.070 |
| Epidemic phase(Established) | 0.019 | 0.128 | 0.877 |
| $\log_{10}$ census value ($N_{i-1}$):Epidemic phase(Epidemic) | -0.009 | 0.034 | 0.791 |
| $\log_{10}$ census value ($N_{i-1}$):Epidemic phase(Established) | -0.035 | 0.052 | 0.493 |

**Supplemental Table 7:** statistical output of a generalized linear mixed model exploring the association between little brown bat colony size in year *i*-1 with the annual colony growth of Indiana bat colonies to year *i*.

| **Model 8 -- Response variable: Annual colony growth rate ($\lambda_i$)**<br>*Generalized linear mixed model with gamma error distribution, log link function, random effect of colony ID (N = 63, groups = 30)*<br>*Model reference level: Epidemic phase(Pre-invasion)* |  |  |  |
| --- | --- | --- | --- |
| *Term* | *Estimate* | *Standard Error* | *P value* |
| Intercept | -0.243 | 0.421 | 0.564 |
| Log<sub>10</sub> MYLU census value ($N_{i-1}$) | 0.115 | 0.176 | 0.514 |
| Epidemic phase(Epidemic) | 0.055 | 0.467 | 0.906 |
| Epidemic phase(Established) | 0.798 | 1.750 | 0.648 |
| Log<sub>10</sub> MYLU census value ($N_{i-1}$):<br>Epidemic phase(Epidemic) | -0.149 | 0.194 | 0.444 |
| Log<sub>10</sub> MYLU census value ($N_{i-1}$):<br>Epidemic phase(Established) | -0.252 | 0.638 | 0.692 |

**Supplemental Table 8:**
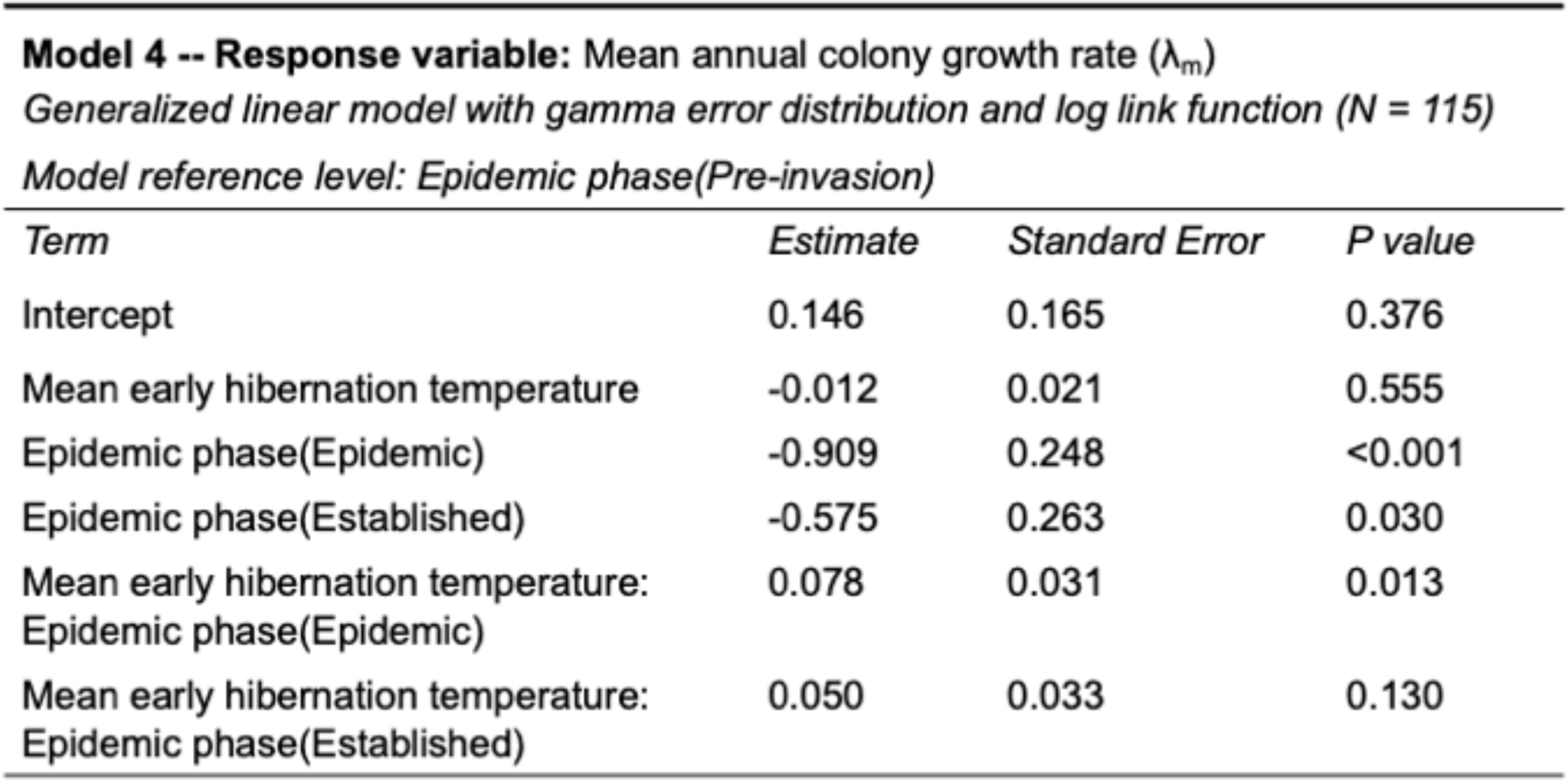
statistical output of a generalized linear model exploring the association between mean early hibernation temperature, epidemic phase, and their interaction and annual colony growth.

**Supplemental Table 9:** statistical output of a generalized linear model exploring the association between mean early hibernation VPD, epidemic phase, and annual colony growth.

| **Model 5 -- Response variable: Mean annual colony growth rate ($\lambda_m$)**<br>*Generalized linear model with gamma error distribution and log link function (N = 81)*<br>*Model reference level: Epidemic phase(Pre-invasion)* |  |  |  |
| --- | --- | --- | --- |
| Term | Estimate | Standard Error | P value |
| Intercept | 0.114 | 0.081 | 0.165 |
| log<sub>10</sub> mean early hibernation VPD | 0.028 | 0.030 | 0.346 |
| Epidemic phase(Epidemic) | -0.292 | 0.118 | 0.015 |
| Epidemic phase(Established) | -0.251 | 0.125 | 0.048 |
| Mean early hibernation VPD: Epidemic phase(Epidemic) | 0.012 | 0.044 | 0.788 |
| Mean early hibernation VPD: Epidemic phase(Established) | -0.034 | 0.046 | 0.469 |

**Supplemental Table 10:**
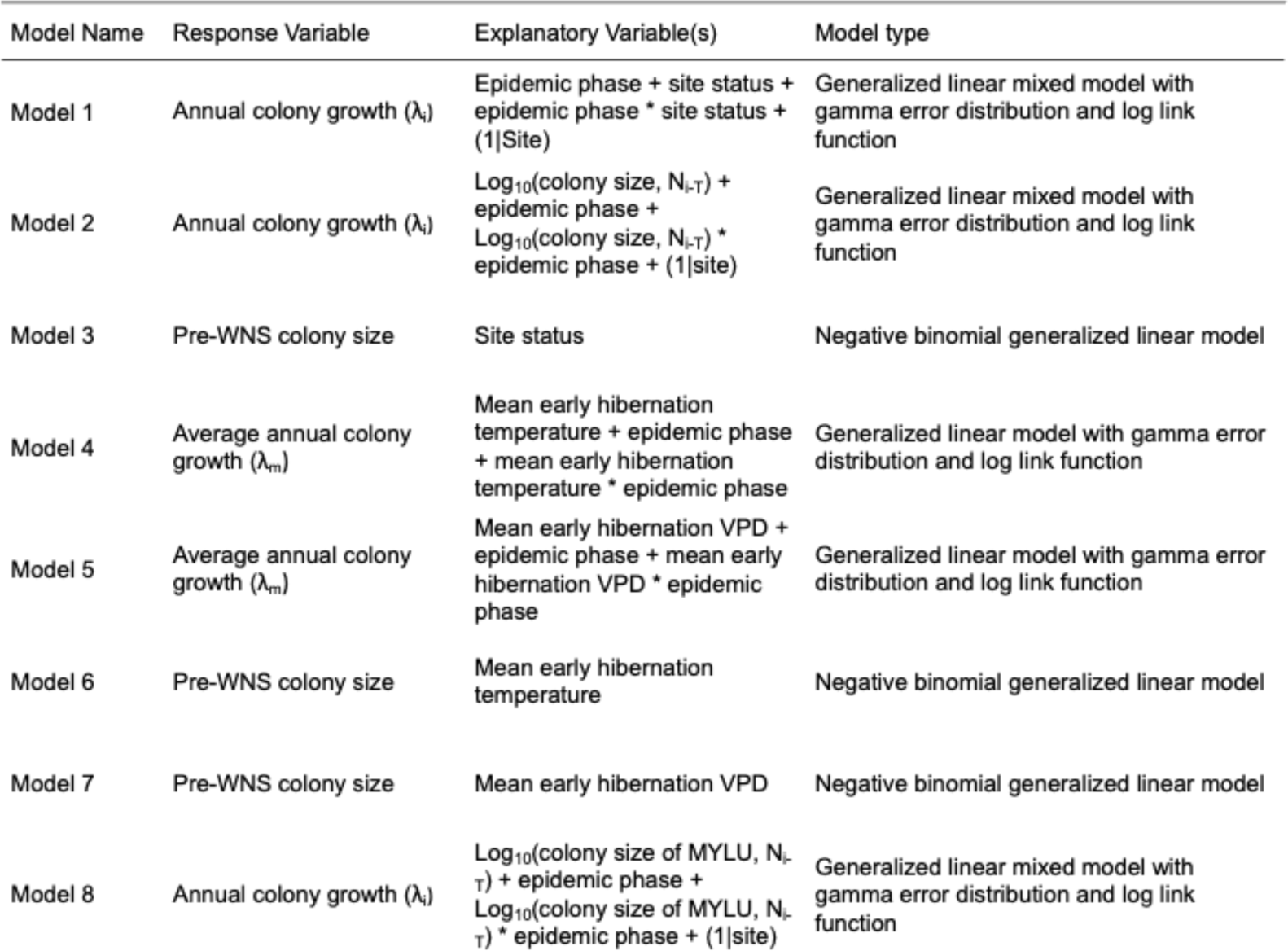
names and descriptions of models reported in this study.

## References

Acevedo, M.A., Beaudrot, L., Meléndez-Ackerman, E.J., Tremblay, R.L., 2020. Local extinction risk under climate change in a neotropical asymmetrically dispersed epiphyte. Journal of Ecology 108, 1553–1564. 10.1111/1365-2745.13361

Altizer, S., Ostfeld, R.S., Johnson, P.T.J., Kutz, S., Harvell, C.D., 2013. Climate change and infectious diseases: from evidence to a predictive framework. Science (1979) 341, 514–519. 10.1126/science.1239401

Anderson, P.K., Cunningham, A.A., Patel, N.G., Morales, F.J., Epstein, P.R., Daszak, P., 2004. Emerging infectious diseases of plants: pathogen pollution, climate change and agrotechnology drivers. Trends Ecol Evol 19, 535–544. 10.1016/J.TREE.2004.07.021

Barrett, R.D.H., Schluter, D., 2008. Adaptation from standing genetic variation. Trends Ecol Evol 23, 38–44. 10.1016/j.tree.2007.09.008

Bell, G., 2013. Evolutionary rescue and the limits of adaptation. Philosophical Transactions of the Royal Society B: Biological Sciences 368, 20120080. 10.1098/rstb.2012.0080

Ben-Hamo, M., Muñoz-Garcia, A., Williams, J.B., Korine, C., Pinshow, B., 2013. Waking to drink: rates of evaporative water loss determine arousal frequency in hibernating bats. Journal of Experimental Biology 216, 573–577. 10.1242/jeb.078790

Blehert, D.S., Hicks, A.C., Behr, M., Meteyer, C.U., Berlowski-Zier, B.M., Buckles, E.L., Coleman, J.T.H., Darling, S.R., Gargas, A., Niver, R., Okoniewski, J.C., Rudd, R.J., Stone, W.B., 2009. Bat white-nose syndrome: An emerging fungal pathogen? Science (1979) 323, 227. 10.1126/science.1163874

Boyles, J.G., Brack, V., McGuire, L.P., 2023. Balancing costs and benefits of managing hibernacula of cavernicolous bats. Mamm Rev. 10.1111/mam.12316

Boyles, J.G., Storm, J.J., Brack, V., Stormf, J.J., 2008. Thermal Benefits of Clustering during Hibernation: A Field Test of Competing Hypotheses on Myotis sodalis. Source: Functional Ecology 22, 632–636. 10.HH/j.1365-2435.2008.01423.x

Boyles, J.G., Willis, C.K.R., 2010. Could localized warm areas inside cold caves reduce mortality of hibernating bats affected by white-nose syndrome? Front Ecol Environ 8, 92–98. 10.1890/080187

Brunner, J.L., Beaty, L., Guitard, A., Russell, D., 2017. Heterogeneities in the infection process drive ranavirus transmission. Ecology 98, 576–582. 10.1002/ecy.1644

Carlson, S.M., Cunningham, C.J., Westley, P.A.H., 2014. Evolutionary rescue in a changing world. Trends Ecol Evol 29, 521–530. 10.1016/J.TREE.2014.06.005

Clawson, R.L., Laval, R.K., La Val, M.L., Caire, W., 1980. CLUSTERING BEHAVIOR OF HIBERNATING MYOTIS SODALIS IN MISSOURI. J Mammal 61, 245–253.

Cohen, J.M., Venesky, M.D., Sauer, E.L., Civitello, D.J., McMahon, T.A., Roznik, E.A., Rohr, J.R., 2017. The thermal mismatch hypothesis explains host susceptibility to an emerging infectious disease. Ecol Lett. 10.1111/ele.12720

Cross, P.C., Heisey, D.M., Scurlock, B.M., Edwards, W.H., Ebinger, M.R., Brennan, A., 2010. Mapping brucellosis increases relative to elk density using hierarchical bayesian models. PLoS One 5, 1–9. 10.1371/journal.pone.0010322

Cryan, P.M., Meteyer, C.U., Blehert, D.S., Lorch, J.M., Reeder, D.M., Turner, G.G., Webb, J., Behr, M., Verant, M., Russell, R.E., Castle, K.T., 2013. Electrolyte depletion in white-nose syndrome bats. J Wildl Dis 49, 398–402. 10.7589/2012-04-121

Cryan, P.M., Meteyer, C.U., Boyles, J.G., Blehert, D.S., 2010. Wing pathology of white-nose syndrome in bats suggests life-threatening disruption of physiology. BMC Biol 8, 1–8. 10.1186/1741-7007-8-135

Cunningham, C.X., Comte, S., McCallum, H., Hamilton, D.G., Hamede, R., Storfer, A., Hollings, T., Ruiz-Aravena, M., Kerlin, D.H., Brook, B.W., Hocking, G., Jones, M.E., 2021. Quantifying 25 years of disease-caused declines in Tasmanian devil populations: host density drives spatial pathogen spread. Ecol Lett 24, 958–969. 10.1111/ele.13703

Daszak, P., Cunningham, A.A., Hyatt, A.D., 2001. Anthropogenic environmental change and the emergence of infectious diseases in wildlife, Acta Tropica.

Daszak, P., Cunningham, A.A., Hyatt, A.D., 2000. Emerging Infectious Diseases of Wildlife. Threats to Biodiversity and Human Health. Science (1979) 287, 443–449. 10.1126/science.287.5452.443

de Castro, F., Bolker, B., 2005. Mechanisms of disease-induced extinction. Ecol Lett 8, 117–126. 10.1111/j.1461-0248.2004.00693.x

Debes, P.V., Gross, R., Vasemägi, A., 2017. Quantitative Genetic Variation in, and Environmental Effects on, Pathogen Resistance and Temperature-Dependent Disease Severity in a Wild Trout. Am Nat 190, 244–265. 10.5061/dryad.12758

Dobony, C.A., Hicks, A.C., Langwig, K.E., von Linden, R.I., Okoniewski, J.C., Rainbolt, R.E., 2011. Little brown myotis persist despite exposure to white-nose syndrome. J Fish Wildl Manag 2, 190–195. 10.3996/022011-JFWM-014

Drees, K.P., Lorch, J.M., Puechmaille, S.J., Parise, K.L., Wibbelt, G., Hoyt, J.R., Sun, K., Jargalsaikhan, A., Dalannast, M., Palmer, J.M., Lindner, D.L., Kilpatrick, A.M., Pearson, T., Keim, P.S., Blehert, D.S., Foster, J.T., 2017. Phylogenetics of a fungal invasion: origins and widespread dispersal of white-nose syndrome. mBio 8, e01941–17. 10.1128/mBio.01941-17

Fisher, M.C., Henk, D.A., Briggs, C.J., Brownstein, J.S., Madoff, L.C., McCraw, S.L., Gurr, S.J., 2012. Emerging fungal threats to animal, plant and ecosystem health. Nature 484, 186–194. 10.1038/nature10947

Frick, W.F., Puechmaille, S.J., Hoyt, J.R., Nickel, B.A., Langwig, K.E., Foster, J.T., Barlow, K.E., Bartonička, T., Feller, D., Haarsma, A.-J., Herzog, C., Horáček, I., van der Kooij, J., Mulkens, B., Petrov, B., Reynolds, R., Rodrigues, L., Stihler, C.W., Turner, G.G., Kilpatrick, A.M., 2015. Disease alters macroecological patterns of North American bats. Global Ecology and Biogeography 24, 741–749. 10.1111/geb.12290

Gamelon, M., Vriend, S.J.G., Engen, S., Adriaensen, F., Dhondt, A.A., Evans, S.R., Matthysen, E., Sheldon, B.C., Sæther, B.E., 2019. Accounting for interspecific competition and age structure in demographic analyses of density dependence improves predictions of fluctuations in population size. Ecol Lett. 10.1111/ele.13237

Gascoigne, J., Berec, L., Gregory, S., Courchamp, F., 2009. Dangerously few liaisons: a review of mate-finding Allee effects. Popul Ecol 51, 355–372. 10.1007/s10144-009-0146-4

Gomulkiewicz, R., Holt, R.D., 1995. When does Evolution by Natural Selection Prevent Extinction?

Grant, E.H.C., Muths, E., Katz, R.A., Canessa, S., Adams, M.J., Ballard, J.R., Berger, L., Briggs, C.J., Coleman, J.T.H., Gray, M.J., Harris, M.C., Harris, R.N., Hossack, B., Huyvaert, K.P., Kolby, J., Lips, K.R., Lovich, R.E., McCallum, H.I., Mendelson, J.R., Nanjappa, P., Olson, D.H., Powers, J.G., Richgels, K.L.D., Russell, R.E., Schmidt, B.R., Spitzen-van der Sluijs, A., Watry, M.K., Woodhams, D.C., White, C.L.A., 2017. Using decision analysis to support proactive management of emerging infectious wildlife diseases. Front Ecol Environ 15, 214–221. 10.1002/fee.1481

Grieneisen, L.E., Brownlee-Bouboulis, S.A., Johnson, J.S., Reeder, D.M., 2015. Sex and hibernaculum temperature predict survivorship in white-nose syndrome affected little brown myotis (*Myotis lucifugus*). R Soc Open Sci 2, 140470. 10.1098/rsos.140470

Grimaudo, A.T., Hoyt, J.R., Yamada, S.A., Herzog, C.J., Bennett, A.B., Langwig, K.E., 2022. Host traits and environment interact to determine persistence of bat populations impacted by white-nose syndrome. Ecol Lett 25, 483–497. 10.1111/ELE.13942

Haddad, N.M., Brudvig, L.A., Clobert, J., Davies, K.F., Gonzalez, A., Holt, R.D., Lovejoy, T.E., Sexton, J.O., Austin, M.P., Collins, C.D., Cook, W.M., Damschen, E.I., Ewers, R.M., Foster, B.L., Jenkins, C.N., King, A.J., Laurance, W.F., Levey, D.J., Margules, C.R., Melbourne, B.A., Nicholls, A.O., Orrock, J.L., Song, D.X., Townshend, J.R., 2015. Habitat fragmentation and its lasting impact on Earth’s ecosystems. Sci Adv 1. 10.1126/sciadv.1500052

Hagman, M., Phillips, B.L., Shine, R., 2009. Fatal attraction: Adaptations to prey on native frogs imperil snakes after invasion of toxic toads. Proceedings of the Royal Society B: Biological Sciences 276, 2813–2818. 10.1098/rspb.2009.0192

Hardin, J.W., Hassell, M.D., 1970. Observation on waking periods and movements of Myotis sodalis during hibernation. J Mammal 51, 829–831.

Heard, G.W., Thomas, C.D., Hodgson, J.A., Scroggie, M.P., Ramsey, D.S.L., Clemann, N., 2015. Refugia and connectivity sustain amphibian metapopulations afflicted by disease. Ecol Lett 18, 853–863. 10.1111/ele.12463

Hodgson, D., McDonald, J.L., Hosken, D.J., 2015. What do you mean, “resilient”? Trends Ecol Evol. 10.1016/j.tree.2015.06.010

Hopkins, S.R., Hoyt, J.R., White, J.P., Kaarakka, H.M., Redell, J.A., DePue, J.E., Scullon, W.H., Kilpatrick, A.M., Langwig, K.E., 2021. Continued preference for suboptimal habitat reduces bat survival with white-nose syndrome. Nat Commun 12, 1–9. 10.1038/s41467-020-20416-5

Hoyt, J.R., Kilpatrick, A.M., Langwig, K.E., 2021. Ecology and impacts of white-nose syndrome on bats. Nat Rev Microbiol 19, 1–15. 10.1038/s41579-020-00493-5

Hoyt, J.R., Langwig, K.E., Okoniewski, J., Frick, W.F., Stone, W.B., Kilpatrick, A.M., 2015. Long-Term Persistence of Pseudogymnoascus destructans, the Causative Agent of White-Nose Syndrome, in the Absence of Bats. Ecohealth 12, 330–333. 10.1007/s10393-014-0981-4

Hoyt, J.R., Langwig, K.E., Sun, K., Parise, K.L., Li, A., Wang, Y., Huang, X., Worledge, L., Miller, H., White, J.P., Kaarakka, H.M., Redell, J.A., Görföl, T., Boldogh, S.A., Fukui, D., Sakuyama, M., Yachimori, S., Sato, A., Dalannast, M., Jargalsaikhan, A., Batbayar, N., Yovel, Y., Amichai, E., Natradze, I., Frick, W.F., Foster, J.T., Feng, J., Kilpatrick, A.M., 2020. Environmental reservoir dynamics predict global infection patterns and population impacts for the fungal disease white-nose syndrome. Proceedings of the National Academy of Sciences 117, 7255–7262. 10.1073/pnas.1914794117/-/DCSupplemental

Huang, Y.H., Joel, H., Küsters, M., Barandongo, Z.R., Cloete, C.C., Hartmann, A., Kamath, P.L., Kilian, J.W., Mfune, J.K.E., Shatumbu, G., Zidon, R., Getz, W.M., Turner, W.C., 2021. Disease or drought: Environmental fluctuations release zebra from a potential pathogen-triggered ecological trap. Proceedings of the Royal Society B: Biological Sciences 288. 10.1098/rspb.2021.0582

Johnson, J.S., Reeder, D.M., McMichael III, J.W., Meierhofer, M.B., F Stern, D.W., Lumadue, S.S., Sigler, L.E., Winters, H.D., Vodzak, M.E., Kurta, A., Kath, J.A., Field, K.A., 2014. Host, Pathogen, and Environmental Characteristics Predict White-Nose Syndrome Mortality in Captive Little Brown Myotis (Myotis lucifugus). PLoS One 9, 112502. 10.1371/journal.pone.0112502

Kilpatrick, A.M., 2011. Globalization, land use, and the invasion of west nile virus. Science (1979) 334, 323–327. 10.1007/sl0869-007-9037-x

Kramer, A.M., Brian, A.E., Ae, D., Liebhold, A.M., Drake, J.M., 2009. The evidence for Allee effects. Popul Ecol 51, 341–354. 10.1007/s10144-009-0152-6

Laggan, N.A., Parise, K.L., White, J.P., Kaarakka, H.M., Redell, J.A., DePue, J.E., Scullon, W.H., Kath, J., Foster, J.T., Kilpatrick, A.M., Langwig, K.E., Hoyt, J.R., 2023. Host infection and disease-induced mortality modify species contributions to the environmental reservoir. Ecology 104. 10.1002/ECY.4147

Lambin, E.F., Tran, A., Vanwambeke, S.O., Linard, C., Soti, V., 2010. Pathogenic landscapes: Interactions between land, people, disease vectors, and their animal hosts. Int J Health Geogr 9. 10.1186/1476-072X-9-54

Lande, R., 1993. Risks of Population Extinction from Demographic and Environmental Stochasticity and Random Catastrophes. Am Nat 142, 911–927.

Lande, R., Engen, S., Saether, B.-E., 2003. Stochastic Population Dynamics in Ecology and Conservation. Oxford University Press.

Langwig, K.E., Frick, W.F., Bried, J.T., Hicks, A.C., Kunz, T.H., Marm Kilpatrick, A., 2012. Sociality, density-dependence and microclimates determine the persistence of populations suffering from a novel fungal disease, white-nose syndrome. Ecol Lett 15, 1050–1057. 10.1111/j.1461-0248.2012.01829.x

Langwig, K.E., Frick, W.F., Hoyt, J.R., Parise, K.L., Drees, K.P., Kunz, T.H., Foster, J.T., Kilpatrick, A.M., 2016. Drivers of variation in species impacts for a multi-host fungal disease of bats. Philosophical Transactions of the Royal Society B: Biological Sciences 371, 20150456. 10.1098/rstb.2015.0456

Langwig, K.E., Frick, W.F., Reynolds, R., Parise, K.L., Drees, K.P., Hoyt, J.R., Cheng, T.L., Kunz, T.H., Foster, J.T.., Kilpatrick, A.M., 2015. Host and pathogen ecology drive the seasonal dynamics of a fungal disease, white-nose syndrome. Proceedings of the Royal Society B 282, 20142335. 10.1098/rspb.2014.2335

Langwig, K.E., Hoyt, J.R., Parise, K.L., Frick, W.F., Foster, J.T., Kilpatrick, A.M., 2017. Resistance in persisting bat populations after white-nose syndrome invasion. Philosophical Transactions of the Royal Society B 372, 20160044. 10.1098/rstb.2016.0044

Leach, C.B., Webb, C.T., Cross, P.C., 2016. When environmentally persistent pathogens transform good habitat into ecological traps. R Soc Open Sci 3. 10.1098/rsos.160051

Leopardi, S., Blake, D., Puechmaille, S.J., 2015. White-Nose Syndrome fungus introduced from Europe to North America. Current Biology 25, R217–R219. 10.1016/j.cub.2015.01.047

Lilley, T.M., Anttila, J., Ruokolainen, L., 2018. Landscape structure and ecology influence the spread of a bat fungal disease. Funct Ecol 32, 2483–2496. 10.1111/1365-2435.13183

Lilley, T.M., Johnson, J.S., Ruokolainen, L., Rogers, E.J., Wilson, C.A., Schell, S.M., Field, K.A., Reeder, D.M., 2016. White-nose syndrome survivors do not exhibit frequent arousals associated with Pseudogymnoascus destructans infection. Front Zool 13, 1–8. 10.1186/s12983-016-0143-3

Lorch, J.M., Knowles, S., Lankton, J.S., Michell, K., Edwards, J.L., Kapfer, J.M., Staffen, R.A., Wild, E.R., Schmidt, K.Z., Ballmann, A.E., Blodgett, D., Farrell, T.M., Glorioso, B.M., Last, L.A., Price, S.J., Schuler, K.L., Smith, C.E., Wellehan, J.F.X., Blehert, D.S., 2016. Snake fungal disease: An emerging threat to wild snakes. Philosophical Transactions of the Royal Society B: Biological Sciences 371, 20150457. 10.1098/rstb.2015.0457

Lorch, J.M., Meteyer, C.U., Behr, M.J., Boyles, J.G., Cryan, P.M., Hicks, A.C., Ballmann, A.E., Coleman, J.T.H., Redell, D.N., Reeder, D.M., Blehert, D.S., 2011. Experimental infection of bats with Geomyces destructans causes white-nose syndrome. Nature 480, 376–379.

Lu, Y., Wu, K., 2011. Effect of relative humidity on population growth of Apolygus lucorum (Heteroptera: Miridae). Appl Entomol Zool 46, 421–427. 10.1007/s13355-011-0058-6

Magnusson, A., Skaug, H., Nielsen, A., Berg, C., Kristensen, K., Maechler, M., van Bentham, K., Bolker, B., Sadat, N., Lüdecke, D., Lenth, R., O’Brien, J., Brooks, M., 2017. Package “glmmTMB” Title Generalized Linear Mixed Models using Template Model Builder. R Package Version 0.2.0.

Mantyka-Pringle, C.S., Visconti, P., Di Marco, M., Martin, T.G., Rondinini, C., Rhodes, J.R., 2015. Climate change modifies risk of global biodiversity loss due to land-cover change. Biol Conserv 187, 103–111. 10.1016/j.biocon.2015.04.016

Marroquin, C.M., Lavine, J.O., Windstam, S.T., 2017. Effect of humidity on development of Pseudogymnoascus destructans, the causal agent of bat white-nose syndrome. Northeast Nat (Steuben) 24, 54–64. 10.1656/045.024.0105

Martin, A.M., Burridge, C.P., Ingram, J., Fraser, T.A., Carver, S., 2018. Invasive pathogen drives host population collapse: Effects of a travelling wave of sarcoptic mange on bare-nosed wombats. Journal of Applied Ecology 55, 331–341. 10.1111/1365-2664.12968

Maslo, B., Fefferman, N.H., 2015. A case study of bats and white-nose syndrome demonstrating how to model population viability with evolutionary effects. Conservation Biology 29, 1176–1185. 10.1111/cobi.12485

Masó, G., Ozgul, A., Fitze, P.S., 2020. Decreased Precipitation Predictability Negatively Affects Population Growth through Differences in Adult Survival. Am Nat 195, 43–55. 10.5061/dryad.349sn3f

Mccallum, H., 2008. Tasmanian devil facial tumour disease: lessons for conservation biology. Trends Ecol Evol 23, 631–637. 10.1016/j.tree.2008.07.001

Mcguire, L.P., Mayberry, H.W., Willis, C.K.R., 2017. White-nose syndrome increases torpid metabolic rate and evaporative water loss in hibernating bats. American Journal of Physiology-Regulatory, Integrative and Comparative Physiology 313, R680–R686. 10.1152/ajpregu.00058.2017.-Fungal

McNew, S.M., Knutie, S.A., Goodman, G.B., Theodosopoulos, A., Saulsberry, A., Yépez R., J., Bush, S.E., Clayton, D.H., 2019. Annual environmental variation influences host tolerance to parasites. Proceedings of the Royal Society B: Biological Sciences 286, 20190049. 10.1098/rspb.2019.0049

Melbourne, B.A., Hastings, A., 2008. Extinction risk depends strongly on factors contributing to stochasticity 454. 10.1038/nature06922

Minnis, A.M., Lindner, D.L., 2013. Phylogenetic evaluation of Geomyces and allies reveals no close relatives of Pseudogymnoascus destructans, comb. nov., in bat hibernacula of eastern North America. Fungal Biol 117, 638–649. 10.1016/j.funbio.2013.07.001

Mordecai, E.A., Caldwell, J.M., Grossman, M.K., Lippi, C.A., Johnson, L.R., Neira, M., Rohr, J.R., Ryan, S.J., Savage, V., Shocket, M.S., Sippy, R., Stewart Ibarra, A.M., Thomas, M.B., Villena, O., 2019. Thermal biology of mosquito-borne disease. Ecol Lett. 10.1111/ele.13335

Mosher, B.A., Bailey, L.L., Muths, E., Huyvaert, K.P., 2018. Host–pathogen metapopulation dynamics suggest high elevation refugia for boreal toads. Ecological Applications 28, 926–937. 10.1002/eap.1699

Ostfeld, R.S., Glass, G.E., Keesing, F., 2005. Spatial epidemiology: An emerging (or re-emerging) discipline. Trends Ecol Evol. 10.1016/j.tree.2005.03.009

Paull, S.H., Song, S., McClure, K.M., Sackett, L.C., Kilpatrick, A.M., Johnson, P.T., 2012. From superspreaders to disease hotspots: linking transmission across hosts and space. Front Ecol Environ 10, 75–82. 10.1890/110111

Puschendorf, R., Hoskin, C.J., Cashins, S.D., Mcdonald, K., Skerratt, L.F., Vanderwal, J., Alford, R.A., 2011. Environmental Refuge from Disease-Driven Amphibian Extinction. Conservation Biology 25, 956–964. 10.1111/j.1523-1739.2011.01728.x

Rachowicz, L.J., Briggs, C.J., 2007. Quantifying the disease transmission function: Effects of density on Batrachochytrium dendrobatidis transmission in the mountain yellow-legged frog Rana muscosa. Journal of Animal Ecology 76, 711–721. 10.1111/j.1365-2656.2007.01256.x

Reeder, D.M., Frank, C.L., Turner, G.G., Meteyer, C.U., Kurta, A., Britzke, E.R., Vodzak, M.E., Darling, S.R., Stihler, C.W., Hicks, A.C., Jacob, R., Grieneisen, L.E., Brownlee, S.A., Muller, L.K., Blehert, D.S., 2012. Frequent arousal from hibernation linked to severity of infection and mortality in bats with white-nose syndrome. PLoS One 7, e38920. 10.1371/journal.pone.0038920

Ripley, B., Venables, B., Bates, D.M., Hornik, K., Gebhardt, A., Firth, D., 2013. Package ‘ MASS.’ Cran r 538, 113–120.

Rogalski, M.A., Gowler, C.D., Shaw, C.L., Hufbauer, R.A., Duffy, M.A., 2017. Human drivers of ecological and evolutionary dynamics in emerging and disappearing infectious disease systems. Philosophical Transactions of the Royal Society B: Biological Sciences 372, 20160043. 10.1098/rstb.2016.0043

Salo, P., Banks, P.B., Dickman, C.R., Korpimäki, E., 2010. Predator manipulation experiments: Impacts on populations of terrestrial vertebrate prey. Ecol Monogr 80, 531–546. 10.1890/09-1260.1

Samuel, M.D., Woodworth, B.L., Atkinson, C.T., Hart, P.J., LaPointe, D.A., 2015. Avian malaria in Hawaiian forest birds: infection and population impacts across species and elevations. Ecosphere 6, 1–21. 10.1890/ES14-00393.1

Scheele, B.C., Skerratt, L.F., Grogan, L.F., Hunter, D.A., Clemann, N., McFadden, M., Newell, D., Hoskin, C.J., Gillespie, G.R., Heard, G.W., Brannelly, L., Roberts, A.A., Berger, L., 2017. After the epidemic: Ongoing declines, stabilizations and recoveries in amphibians afflicted by chytridiomycosis. Biol Conserv 206, 37–46. 10.1016/j.biocon.2016.12.010

Şekercioĝlu, çaĝan H., Primack, R.B., Wormworth, J., 2012. The effects of climate change on tropical birds. Biol Conserv. 10.1016/j.biocon.2011.10.019

Skerratt, L.F., Berger, L., Speare, R., Cashins, S., McDonald, K.R., Phillott, A.D., Hines, H.B., Kenyon, N., 2007. Spread of chytridiomycosis has caused the rapid global decline and extinction of frogs. Ecohealth 4, 125–134. 10.1007/s10393-007-0093-5

Smith, M.J., Telfer, S., Kallio, E.R., Burthe, S., Cook, A.R., Lambin, X., Begon, M., 2009. Host-pathogen time series data in wildlife support a transmission function between density and frequency dependence. Proc Natl Acad Sci U S A 106, 7905–7909. 10.1073/pnas.0809145106

Spitzen-Van Der Sluijs, A., Canessa, S., Martel, A., Pasmans, F., 2017. Fragile coexistence of a global chytrid pathogen with amphibian populations is mediated by environment and demography. Proceedings of the Royal Society B: Biological Sciences 284, 20171444. 10.1098/rspb.2017.1444

Stephens, P.A., Sutherland, W.J., Freckleton, R.P., 1999. What Is the Allee Effect? Oikos 87, 185–190.

Storm, D.J., Samuel, M.D., Rolley, R.E., Shelton, P., Keuler, N.S., Richards, B.J., Van Deelen, T.R., 2013. Deer density and disease prevalence influence transmission of chronic wasting disease in white-tailed deer. Ecosphere 4. 10.1890/ES12-00141.1

Sutherland, W.J., Pullin, A.S., Dolman, P.M., Knight, T.M., 2004. The need for evidence-based conservation. Trends Ecol Evol 19, 305–308. 10.1016/j.tree.2004.03.018

Thogmartin, W.E., King, R.A., Mckann, P.C., Szymanski, J.A., Pruitt, L., 2012. Population-level impact of white-nose syndrome on the endangered Indiana bat. J Mammal 93, 1086–1098. 10.1644/11-MAMM-A-355.1

Thomas, D.W., Cloutier, D., 1992. Evaporative Water Loss by Hibernating Little Brown Bats, Myotis lucifugus. Physiol Zool 65, 443–456.

Thomas, D.W., Dorais, M., Bergeron, J., 1990. Winter Energy Budgets and Cost of Arousals for Hibernating Little Brown Bats, Myotis lucifugus. J Mammal 71, 475–479.

Thomson, C.E., 1982. Myotis sodalis. Mammalian Species 1–5. 10.2307/3504013

Turner, J.M., Warnecke, L., Wilcox, A., Baloun, D., Bollinger, T.K., Misra, V., Willis, C.K.R., 2014. Conspecific disturbance contributes to altered hibernation patterns in bats with white-nose syndrome. Physiol Behav 140, 71–78. 10.1016/j.physbeh.2014.12.013

Van Dyck, H., Bonte, D., Puls, R., Gotthard, K., Maes, D., 2015. The lost generation hypothesis: Could climate change drive ectotherms into a developmental trap? Oikos 124, 54–61. 10.1111/oik.02066

Van Riper III, C., Van Riper, S.G., Goff, M.L., Laird, M., 1986. The epizootiology and ecological significance of malaria in Hawaiian land birds. Ecol Monogr 56, 327–344.

Verant, M.L., Boyles, J.G., Waldrep Jr., W., Wibbelt, G., Blehert, D.S., 2012. Temperature-dependent growth of *Geomyces destructans*, the fungus that causes bat white-nose syndrome. PLoS One 7, e46280. 10.1371/journal.pone.0046280

Verant, M.L., Meteyer, C.U., Speakman, J.R., Cryan, P.M., Lorch, J.M., Blehert, D.S., 2014. White-nose syndrome initiates a cascade of physiologic disturbances in the hibernating bat host. BMC Physiol 14. 10.1186/s12899-014-0010-4

Warnecke, L., Turner, J.M., Bollinger, T.K., Lorch, J.M., Misra, V., Cryan, P.M., Wibbelt, G., Blehert, D.S., Willis, C.K.R., 2012. Inoculation of bats with European *Geomyces destructans* supports the novel pathogen hypothesis for the origin of white-nose syndrome. Proceedings of the National Academy of Sciences 109, 6999–7003. 10.1073/pnas.1200374109

Warnecke, L., Turner, J.M., Bollinger, T.K., Misra, V., Cryan, P.M., Blehert, D.S., Wibbelt, G., Willis, C.K.R., 2013. Pathophysiology of white-nose syndrome in bats: a mechanistic model linking wing damage to mortality. Biol Lett 9, 20130177. 10.1098/rsbl.2013.0177

Webb, P.I., Speakman, J.R., Racey, P.A., 1995. Evaporative water loss in two sympatric species of vespertilionid bat, Plecotus auritus and Myotis daubentoni: relation to foraging mode and implications for roost site selection. J Zool 235, 269–278. 10.1111/j.1469-7998.1995.tb05143.x

Woodworth, B.K., Wheelwright, N.T., Newman, A.E., Schaub, M., Norris, D.R., 2017. Winter temperatures limit population growth rate of a migratory songbird. Nat Commun 8. 10.1038/ncomms14812

